# Transcriptomic and epigenetic responses shed light on soybean resistance to *Phytophthora sansomeana*

**DOI:** 10.1101/2024.01.27.577583

**Authors:** Gwonjin Lee, Charlotte N. DiBiase, Beibei Liu, Tong Li, Austin G. McCoy, Martin I. Chilvers, Lianjun Sun, Dechun Wang, Feng Lin, Meixia Zhao

**Affiliations:** Department of Microbiology and Cell Science, University of Florida, Gainesville, FL 32611, USA; Department of Biology, Miami University, Oxford, OH 45056, USA; College of Agronomy and Biotechnology, China Agricultural University, Beijing 100193, China; Department of Plant, Soil and Microbial Sciences, Michigan State University, East Lansing, MI 48824, USA; Fisher Delta Research, Extension, and Education Center, Division of Plant Sciences and Technology, University of Missouri, Portageville, MO 63873, USA

**Author notes:** Corresponding authors Feng Lin and Meixia Zhao.

**Keywords:** *Phytophthora sansomeana*, soybean, transcriptomics, long non-coding RNAs, DNA methylation

## Abstract

Phytophthora root rot caused by oomycete pathogens in the Phytophthora genus poses a significant threat to soybean productivity. While resistance mechanisms against *Phytophthora sojae* have been extensively studied in soybean, the molecular basis underlying immune responses to *Phytophthora sansomeana* remains unclear. We investigated transcriptomic and epigenetic responses of two resistant (Colfax and NE2701) and two susceptible (Williams 82 and Senaki) soybean lines at four time points (2, 4, 8, and 16 hours post inoculation, hpi) after *P. sansomeana* inoculation. Comparative transcriptomic analyses revealed a greater number of differentially expressed genes (DEGs) upon pathogen inoculation in resistant lines, particularly at 8 and 16 hpi, predominantly associated with ethylene and reactive oxygen species-mediated defense pathways. Moreover, DE transposons were predominantly up-regulated after inoculation and enriched near genes in Colfax. A long non-coding RNA (lncRNA) within the mapped region of the resistance gene exhibited exclusive up-regulation in the resistant lines after inoculation, potentially regulating two flanking *LURP-one-related* genes. Furthermore, DNA methylation analysis revealed increased CHH methylation levels in lncRNAs after inoculation, with delayed responses in Colfax compared to Williams 82. Overall, our results provide comprehensive insights into soybean responses to *P. sansomeana*, highlighting potential roles of lncRNAs and epigenetic regulation in plant defense.

## Introduction

Plants are constantly challenged by diverse pathogens, including bacteria, fungi, viruses, and oomycetes. To counter these biotic stresses, plants have evolved diverse defense mechanisms (J. D. G. Jones & J. L. Dangl, 2006). Cell surface receptor proteins called pattern recognition receptors in plants recognize pathogen-associated molecular patterns (PAMPs), eliciting PAMP-triggered immunity (PTI) (Boller & He, 2009). Additionally, plants have intracellular nucleotide binding site-leucine rich repeat receptors (NBS-LRRs or NLRs) that can recognize pathogen effectors, leading to effector-trigger immunity (ETI) (Dodds & Rathjen, 2010; Ngou et al., 2022). Both PTI and ETI could facilitate rapid cellular responses, such as calcium influx, reprogramming of the expression of defense-responsive genes (Amorim et al., 2017; Thirugnanasambantham et al., 2015), and activation of mitogen-activated protein kinases (MAPKs) that coordinate immune responses and modulate other defense-related genes (Meng & Zhang, 2013). Plant defense against pathogens also involves the production of defense phytohormones, including ethylene (ET), salicylic acid (SA), jasmonic acid (JA), brassinosteroids (BR), and auxin. These hormones collaboratively work to regulate immune responses in plants by serving as the primary molecules in the induced defense signaling network (Bürger & Chory, 2019; Pieterse et al., 2009). Additionally, in response to pathogenic attacks, plants produce specific compounds such as phytoalexins (Ahuja et al., 2012; Hammerschmidt, 1999) and reactive oxygen species (ROS) (Mohammadi et al., 2021; Tyagi et al., 2022). Transcriptional changes for the coordination of these multiple pathways collectively contribute to the complex defense system that enables plants to effectively counteract pathogen challenges.

In addition to transcriptional responses in protein encoding genes, plants react to environmental stresses by rapidly altering the transcription and activity of other genetic elements, such as long non-coding RNAs (lncRNAs) and transposable elements (TEs) (Guo et al., 2021; Hou et al., 2019; Klein & Anderson, 2022). LncRNAs are a class of RNAs longer than 200 nucleotides that lack protein-coding potential (Chekanova, 2015; Mattick et al., 2023). Despite their inability to code for proteins, lncRNAs play crucial roles in regulating gene expression both in *cis* and in *trans* through mechanisms such as transcriptional or post-transcriptional regulation, chromatin modification, RNA processing and stability, and scaffold interactions (Sharma et al., 2022; Zhang et al., 2020). While high-throughput RNA sequencing technologies have enabled the identification of a significant number of lncRNAs in plants associated with defense responses against fungal, viral, and bacterial infections (Di et al., 2014; Joshi et al., 2016; Seo et al., 2017; Sun et al., 2020; Wang et al., 2015; Wang et al., 2017; Xin et al., 2011; Yu et al., 2020; Zhang et al., 2013; Zhu et al., 2014), functional characterization of these lncRNAs remains limited. Particularly noteworthy is the scarcity of studies that have explored the specific roles of lncRNAs in conferring resistance to pathogenic oomycetes. A primary focus of lncRNAs to oomycetes has been on *Phytophthora infestans*, the causal pathogen responsible for late blight in tomato (Cui et al., 2020; Cui et al., 2017; Su et al., 2023). These identified lncRNAs modulate a range of genes that control the activation of multiple phytohormones and the accumulation of ROS, thereby enhancing the plants’ resistance to the pathogen.

DNA methylation is a fundamental epigenetic process that regulates transcription activity, transposon mobility, and chromatin stability in response to both abiotic and biotic stresses (Arora et al., 2022; Dowen et al., 2012; Hewezi et al., 2018). In plants, DNA methylation commonly occurs in three cytosine sequence contexts CG, CHG, and CHH (where H represents A, T, or C) (Law & Jacobsen, 2010; Matzke & Mosher, 2014). The alternation of methylation levels in genes and TEs plays a pivotal role in orchestrating the dynamic rewiring of plant genomes during stress responses to pathogen virulence (Cambiagno et al., 2018; Liu & Zhao, 2023; Zervudacki et al., 2018). For instance, a study investigating DNA methylation variants associated with soybean cyst nematode parasitism reveals distinct methylation patterns across three cytosine contexts (Rambani et al., 2020). Furthermore, methylation changes have been noted in lncRNAs of several species (Ding et al., 2012; He et al., 2014; Li et al., 2021; Zhao et al., 2016). A notable example is the rice lncRNA LDMAR, which experiences increased methylation in its promoter region due to a point mutation, resulting in abnormal pollen development (Ding et al., 2012). Nevertheless, the specific mechanisms underlying the methylation changes of lncRNAs in the context of plant immunity remains largely unexplored.

Phytophthora root rot (PRR) is one of the most destructive diseases in soybean (*Glycine max*), causing substantial soybean yield losses worldwide (Allen et al., 2017; Kaufmann & Gerdemann, 1958; Sahoo et al., 2021; Sandhu et al., 2005). Historically, this disease has been attributed to the soil-borne hemibiotrophic oomycete pathogen, *Phytophthora sojae*, which primarily infects soybean (Dorrance et al., 2008; Malvick & Grunden, 2004; TYLER, 2007). The resistance to *P. sojae* (*Rps*) genes have been effective in controlling PRR, with over 40 *Rps* genes/alleles identified as crucial regulators modulating diverse pathways against *P. sojae* (Anderson & Buzzell, 1992; Burnham et al., 2003; Lin et al., 2014; Polzin et al., 1994; Sahoo et al., 2021; Sandhu et al., 2005; Zhou et al., 2022). Many of these *Rps* genes belong to the NBS-LRR family, which can directly or indirectly recognize the corresponding effectors from a variety of pathogens (Ngou et al., 2022; Wu et al., 2018). In 2009, *Phytophthora sansomeana* was differentiated from the *Phytophthora megasperma* complex as a causal agent of root rot across a broader range of hosts, including soybean, carrot, pea, white clover, gerbera, and maize (Detranaltes et al., 2022; Hansen et al., 2009; Lin et al., 2021; Rojas et al., 2017; Zelaya-Molina et al., 2010). The prevalence of *P. sansomeana* in soybean, particularly in North America and Northeast Asia, has led to significant agricultural losses (Alejandro Rojas et al., 2017; Lin et al., 2021; Malvick & Grunden, 2004; Rahman et al., 2015; Tang et al., 2010). In a recent study on the pathogenicity of oomycete species, it was determined that *P. sansomeana* has a more virulent impact on root reduction in soybean seedlings compared to *P. sojae* (Alejandro Rojas et al., 2017). However, unlike *P*. *sojae*, no resistant genes have been identified in soybean for *P. sansomeana*, leaving the molecular and physiological mechanisms underlying defense or stress responses to this pathogen poorly understood. Therefore, further exploration into the genetic control of immune responses to *P. sansomeana* becomes imperative to understand its pathogenicity and interactions with soybean for fortifying crop defenses.

Here, we conducted comprehensive analyses of transcriptome landscapes, including an in-depth investigation of differential transcriptions of genes, TEs, and lncRNAs, along with DNA methylation patterns in response to *P. sansomeana* to unravel the genetic and epigenetic basis of defense mechanisms in the resistant soybean lines by comparison with the susceptible lines. Our comparative analysis revealed contrasting transcriptomic and epigenetic responses between the resistant and susceptible lines. Specially, we observed the specific expression of a number of differentially expressed genes (DEGs) linked to ET-mediated defense responses, along with ROS generation with increased hydrogen peroxide levels within the resistant lines. Furthermore, the DE TEs were prominently up-regulated after inoculation, particularly in proximity to genes in Colfax. We also identified a DE lncRNA within the mapped region of the resistance gene in Colfax that exhibited significant increase in expression exclusively within the resistant lines following pathogen inoculation. This lncRNA holds the potential to regulate adjacent *LURP-one-related* (*LOR*) genes, known for their involvement in defense-related mechanisms against pathogenic oomycetes in Arabidopsis (Baig, 2018; Knoth & Eulgem, 2008). Additionally, our DNA methylation analysis revealed increased CHH methylation levels in lncRNAs after inoculation, at a later time point in the resistant lines compared to an earlier time point in the susceptible lines, suggesting distinctive epigenetic responses of these different soybean lines. Together, our study provides valuable insights into the molecular and physiological mechanisms underlying the defense responses of soybean to the newly recognized pathogen *P. sansomeana*.

## Materials and Methods

### Plant growth, inoculation, and sample collection

Four soybean lines, Colfax, NE2701, Senaki, and Williams 82 were grown in the greenhouse at Michigan State University. For each of the four biological replicates, we divided the seedlings from each line into two groups: one group was inoculated with *P. sansomeana*, while the other group was mock-inoculated without the pathogen. The cultivation of *P. sansomeana* followed established protocols typically used for studying *P. sojae* (Dorrance et al., 2008). In each replicate, ten seedlings from each line were challenged with the *P. sansomeana* isolate *MPS17-22* using the standard hypocotyl inoculation method (Lin et al., 2021). Tissues were collected from each line and condition at four designated time points: 2 hours post inoculation (hpi), 4 hpi, 8 hpi, and 16 hpi. Specifically, stem tissues were collected from 7-8 seedlings in each replicate by excising a 2-3 cm segment from the wounded site. These collected tissues were rapidly frozen using liquid nitrogen and subsequently stored at −80°C. Additionally, the remaining seedlings were retained to closely monitor the progression of symptoms and assess survival rates for a period up to one week post inoculation.

### RNA sequencing and data analysis

Total RNA was extracted from approximately 100 mg of frozen stem tissues for each sample using the Qiagen RNeasy Plant Mini-Kit (Qiagen, Valencia, CA) according to the manufacturer’s instructions. Subsequently, library preparation and RNA sequencing were conducted at Novogene (Sacramento, CA) using the Illumina HiSeq 4000 and NovaSeq 6000 platform. The sequencing yielded a total of 30 to 47 million 150 bp read pairs per sample, which were used for further transcriptome analyses.

The quality of the sequenced reads was assessed using FastQC v0.11.7 (http://www.bioinformatics.babraham.ac.uk/projects/fastqc/). The raw reads from the RNA-seq data were trimmed using Trimmomatic v0.36 to remove adapters and low-quality reads (Bolger et al., 2014). The trimmed reads were then aligned to the soybean reference genome, Williams 82 genome assembly v4.0 (Wm82.a4), using HISAT v2.2.1 (Kim et al., 2015; Valliyodan et al., 2019). Additionally, uniquely mapped reads were retained, and reads with multiple matches were maintained by opting the shortest alignments or selecting a single random alignment for each instance of palindromic multiple mapping using selected scripts from COMEX v2.1 (Lee et al., 2021; Pietzenuk et al., 2016). The number of reads per gene for each sample was quantified using HTSeq v0.11.2 (Putri et al., 2022).

Differentially expressed genes (DEGs) between the control and inoculated samples were identified, and principal component analysis (PCA) of count data was conducted to assess data quality and sample distance using the R package *DESeq2* v1.12.3 (Love et al., 2014). Significant DEGs were identified using adjusted *P*-values from Wald-Test with FDR correction (Benjamini & Hochberg, 1995). Normalized counts obtained from *DESeq2* were used to compare transcript levels for genes across samples. Furthermore, co-expressed gene clusters were identified for each line using Clust v1.18.1 (Abu-Jamous & Kelly, 2018). One or two clusters per line, displaying similar patterns such as clusters up-regulated after inoculation, were selected to identify co-expressed genes specific to the resistant or susceptible lines.

To determine biological categories of DEGs or genes selected in cluster analysis, gene ontology (GO) enrichment analysis was conducted using g:GOst functional profiling in g:Profiler (Raudvere et al., 2019). This analysis employed a hypergeometric test to identify significantly overrepresented GO terms (adjusted *P* < 0.01). In addition, heatmaps depicting relative expression levels per gene for different pathways related to the response to *P. sansomeana* were generated. We manually curated a custom annotated list of genes potentially involved in response pathways, including signaling or synthesis of ET, JA, SA, BR, and MAPK. Normalized counts of these genes were used for measuring transcript levels, and gene clustering based on Euclidean distance was performed using the R package *pheatmap*.

### Analysis of transposable elements

To explore the transcriptional activities of TEs, the mapped and corrected reads obtained from RNA-seq were counted using HTSeq v0.11.2 based on the TE annotation of the Wm82.a4 genome (Valliyodan et al., 2019). We focused on six superfamilies of DNA transposons and four superfamilies of retrotransposons (see results). To compare the overall transcript levels of each TE superfamily across different lines and conditions, we used normalized counts divided by the length of each TE (NCPK; normalized counts per kilobase of TE length) as described in (Lee et al., 2021). Additionally, we counted the numbers and proportions of transcribed TEs (NCPK > 0.5) per superfamily per sample. Moreover, differentially transcribed TEs were identified using the R package *DESeq2* v1.12.3, the same as the method employed in the DEG analysis. Subsequently, we calculated the proportions of TE superfamilies among the DE TEs and compared them to the genome-wide levels. To determine the genomic locations of DE TEs, we examined whether they were located within or in close proximity to genes, as well as in pericentromeric regions and chromosomal arms based on annotations for TEs and genes using the *intersect* and *window* functions in bedtools v2.30.0 (Quinlan & Hall, 2010).

### LncRNA identification and analysis

To identify lncRNAs, the transcriptomic data was *de novo* assembled following the Cufflinks v2.2.1.1 workflow (Trapnell et al., 2010). To achieve this, trimmed reads from the RNA-seq data were mapped to the soybean reference genome, Wm82.a4, using HISAT 2.2.1, with the inclusion of the Cufflinks option (--dta-cufflinks). The resultant mapped reads were used for transcript assembly for each sample through Cufflinks. These assembled transcripts were then merged to generate the final transcriptome assembly using the Cuffmerge function.

LncRNAs (> 200 bp) were identified from the assembled and merged transcripts using Evolinc-I v1.7.5 (Nelson et al., 2017). This identification process categorized lncRNAs into three types: long intergenic non-coding RNAs (lincRNAs), sense-overlapping lncRNAs (SOTs), and antisense (AOTs). To differentiate TE-containing lncRNAs from non-TE lincRNAs, we incorporated TE sequence data obtained using gff3toolkit v2.0.3 (Chen et al., 2019), based on the manually refined annotation of the repeat-masked assembly for the Wm82.a4 genome with the sequence similarity bit score > 200 and E-value < 1E-20.

Furthermore, transcription patterns of four transcript types, including genes, non-TE lincRNAs, TE-containing lincRNAs, and TEs, were explored using NCPK as the measure of transcript levels. First, the proportion of transcribed transcripts (NCPK > 0.5) across different conditions and lines using all samples was calculated for each transcript type. Subsequently, transcript levels of these transcripts were compared using log(NCPK + 1).

For differentially transcribed lincRNAs upon inoculation, a method similar to the DEG analysis was pursued. DE lincRNAs were separately analyzed for non-TE lincRNAs and TE-containing lincRNAs, using the R package *DESeq2* v1.12.3 (as detailed above). Extracted normalized counts were used to examine the transcript levels of DE lincRNAs across different lines and conditions. We further investigated target genes by analyzing DE lincRNAs that were exclusively identified in either resistant or susceptible lines. To uncover potential *trans*-target genes, correlation analysis was employed between the transcript levels of DEGs and DE lincRNAs. DEGs with correlation coefficients (*R*) greater than 0.85 or less than −0.85, alongside a *P*-value < 0.001, were considered potential *trans*-targets. Additionally, for potential *cis*-targets, genes or TEs located within 5 kb upstream and downstream of DE lincRNAs were extracted using bedtools v2.30.0.

### qRT-PCR validation

Total RNA for qRT-PCR was extracted from a separate biological replicate, distinct from the samples used for transcriptome sequencing, using the RNeasy Plant Mini with DNase I treatment according to the manufacturer’s instructions (Qiagen, Valencia, CA). One μg of RNA was used for a 20-µL first-strand cDNA synthesis reaction using High Capacity cDNA Reverse Transcription Kit (Thermo Fisher Scientific, Waltham, MA) following the manufacturer’s guidelines. For each 10 µL of qPCR reaction, 2 µL of a 1/10 dilution of the synthesized cDNA was mixed with PowerTrack SYBR Green Master Mix (Thermo Fisher Scientific, Waltham, MA) and primer pairs. The thermal cycling involved initial denaturation at 95°C for 2 min, 40 cycles at 95°C for 15 sec and 60°C for 1 min, followed by final elongation at 60°C for 10 min, with a dissociation step. Quantification of relative transcript levels (RTL) involved normalizing the transcript levels of genes of interest to the respective transcript level of the housekeeping gene *Cons4* that had lowest variation in transcript levels among housekeeping genes across different samples in our RNA sequencing dataset.

### Determination of hydrogen peroxide concentration

Methods for the measurement of hydrogen peroxide (H_2_O_2_) concentration followed procedures previously described (Alexieva et al., 2001). In brief, tissue samples were ground in liquid nitrogen and mixed with 0.5 ml of 0.1% trichloroacetic acid. Following centrifugation, 0.25 ml of 0.1 M phosphate buffer (pH 7.0) and 1 ml of 1 M KI were added to 0.25 ml of the resulting supernatant. The absorbance was then measured at 390 nm using the spectrophotometer GENESYS 20 (Thermo Fisher Scientific, Waltham, MA) after the reaction rested for 1h in darkness. The quantification of H_2_O_2_ was determined by a standard curve prepared using known concentrations of H_2_O_2_.

### Analysis of whole genome bisulfite sequencing data

DNA was isolated from the same tissues of two of the four soybean lines, Colfax and Willaims 82, at two time points (4 and 16 hpi) using the modified CTAB method. Library construction and subsequent sequencing were performed at Novogene. The raw reads were quality controlled by FastQC and trimmed using Trimmomatic (Bolger et al., 2014). The resulting clean reads were then mapped to the soybean reference genome v4.0 (Wm82.a4) using Bismark with the following parameters (-I 50, -N 1) (Krueger & Andrews, 2011; Valliyodan et al., 2019). To eliminate PCR duplicates, the deduplication package under Bismark was utilized. Additional packages under Bismark, including Bismark methylation extractor, bismark2bedGraph and coverage2cytosine, were employed to extract methylated cytosines and count methylated and unmethylated reads following our previous research (Yin et al., 2022; Zhao et al., 2021).

Methylated levels for each cytosine contexts (CG, CHG, and CHH) were calculated by dividing the number of methylated reads by the total number of methylated and unmethylated reads (Schultz et al., 2012). The average methylation level for each cytosine context per transcript of non-TE lincRNAs was calculated for each line and condition using the *map* function in bedtools v2.30.0. Additionally, we examined the distribution of methylation proportions across three genomic regions: transcript bodies, 2 kb upstream, and 2 kb downstream of the lincRNAs. Mean methylation levels were calculated in 40 windows per region to assess the overall methylation patterns across lincRNA bodies and their flanking regions.

## Results

### Two soybean lines, Colfax and NE2701, are resistant to *P. sansomeana*

We previously screened approximately 500 soybean lines to identify resistance to *P. sansomeana* and discovered two lines, Colfax and NE2701, which exhibited resistance to the pathogen in both field and greenhouse conditions. Alongside these two resistant lines, we included two susceptible lines, Senaki and Williams 82, the latter of which serves as the reference genome of soybean, for transcriptomic analyses. Four biological replicates from each line were planted under greenhouse conditions. The seedlings from each line were then divided into two groups: one group was inoculated with *P. sansomeana*, while the other group was mock-inoculated without the pathogen. We observed stem rot in the majority of Senaki and Willams 82 plants three days after pathogen inoculation, while a few Colfax and NE2701 plants exhibited the symptom (Figure S1). Additionally, we evaluated the resistance of the four soybean lines by measuring their survival rates to *P. sansomeana* seven days post mock (control) and pathogen inoculation (inoculated) to validate resistance or susceptibility of the four selected soybean lines to the pathogen. The survival rates for all four lines were consistently 100% after mock inoculation (Figure 1). The average survival rates for both resistant lines, Colfax and NE2701, showed no significant difference between mock and pathogen inoculation. However, the two susceptible lines, Senaki and Williams 82, displayed significant susceptibility to *P. sansomeana* after pathogen inoculation, when compared to mock inoculation (Figure 1; ANOVA, followed by Tukey’s HSD test; *P* < 0.05). The average survival rates of Senaki and Williams 82 upon pathogen inoculation were approximately 70% and 80% lower than those after mock inoculation, respectively. These results demonstrate that Colfax and NE2701 exhibit resistance to *P. sansomeana*, while Senaki and Williams 82 are susceptible to the pathogen.

**Figure 1.**
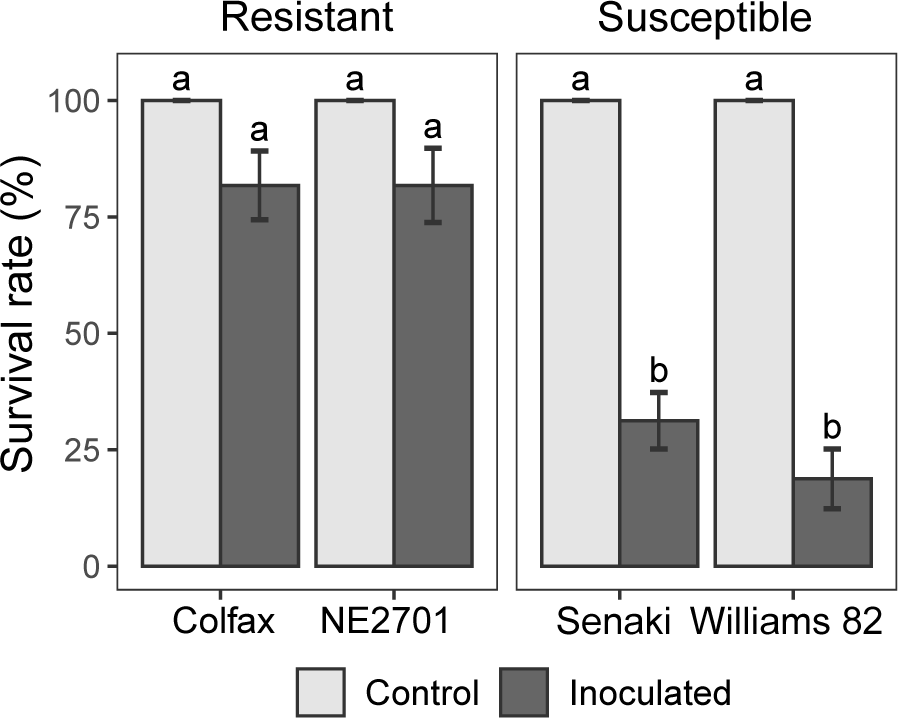
Survival rates of four soybean lines after mock (control) and *P. sansomeana* inoculation (inoculated). Survival seedlings were counted seven days post inoculation. The results are presented as mean ± SE (*n* = 16 per line per treatment). Different characters were used to denote statistically significant differences between means based on two-way ANOVA, followed by Tukey’s HSD test (*P* < 0.05).

### Transcriptomic responses to pathogen inoculation reveal contrasting patterns in resistant and susceptible soybean lines

To dissect the molecular responses to the pathogen in the resistant and susceptible lines, we conducted a comprehensive analysis of transcriptomic changes following inoculation. A total of 64 soybean samples, including four different lines, four time points (2, 4, 8, and 16 hours post inoculation, hpi), and two treatments (pathogen vs. mock), each with two biological replicates, were used for transcriptome analyses. The principal component analysis (PCA) of the transcriptome revealed that the primary source of variation (PC1; explained 54% of total variance) was attributed to four different time points, while the second component, PC2 (14%), was associated with inoculation types (Figure S2).

Next, we identified differentially expressed genes (DEGs) in response to *P. sansomeana* inoculation. Upon pathogen inoculation, the transcriptomic differences in both the resistant lines comprised a considerably greater number of DEGs than those in the susceptible lines (Figure 2a; Dataset S1). In the resistant line, Colfax, a total of 5,058 genes were significantly up-regulated, and 3,120 genes were down-regulated across the four time points in response to the pathogen. Similarly, in the other resistant line, NE2701, 2,611 genes were up-regulated, and 1,364 genes were down-regulated upon pathogen inoculation. In contrast, the susceptible lines, Senaki and Williams 82, exhibited a relatively smaller number of DEGs following inoculation. Senaki showed 1069 up-regulated and 158 down-regulated genes, while Williams 82 had 603 up-regulated and 256 down-regulated genes (Figure 2a).

**Figure 2.**
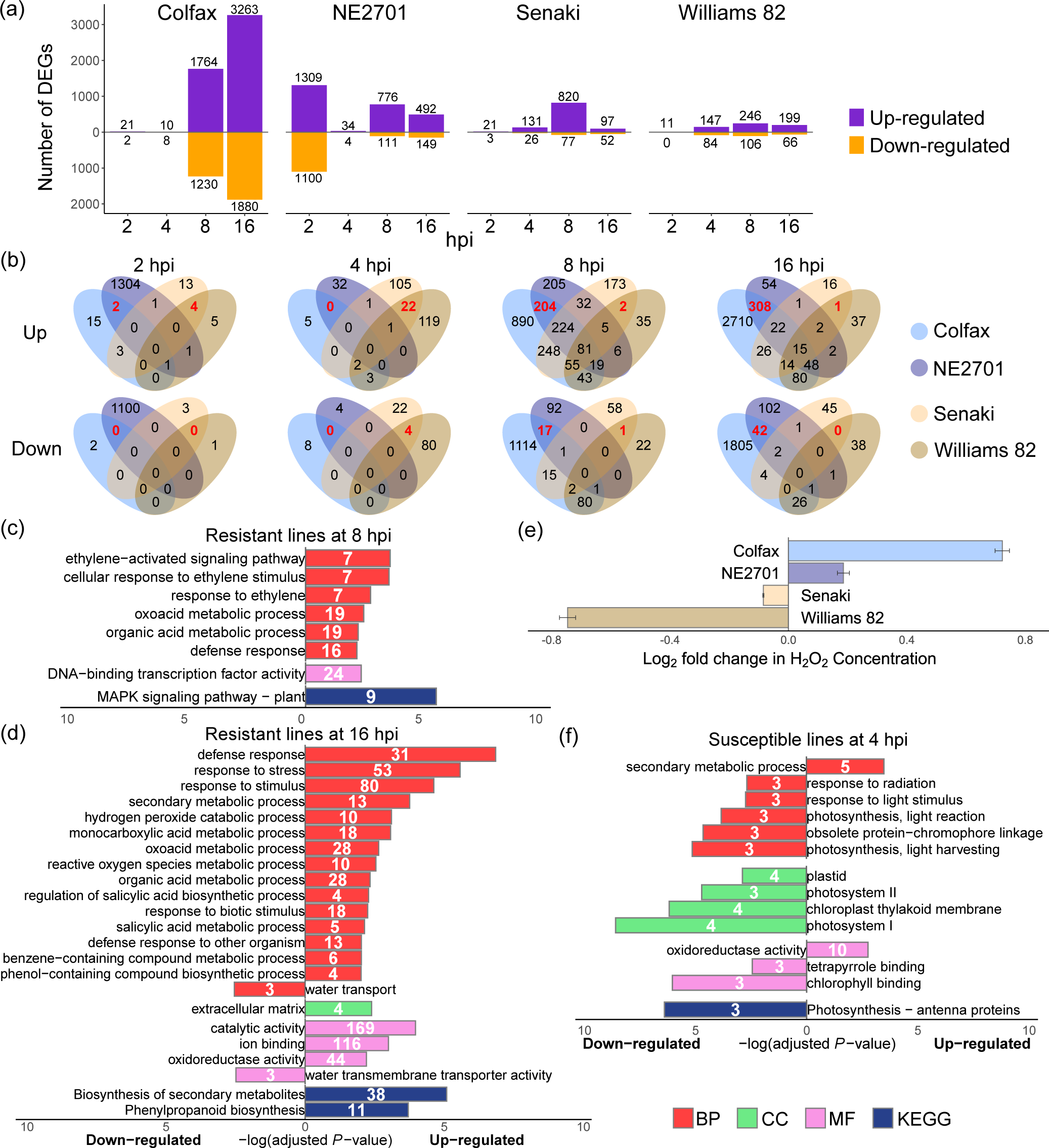
Transcriptomic responses to pathogen inoculation reveal contrasting patterns in the resistant and susceptible soybean lines. (a) The number of significant DEGs for four lines (adjusted *P* < 0.05) at four different time points (2, 4, 8, and 16 hours post inoculation, hpi). (b) Shared and unique up-and down-regulated DEGs at each time point. The upper row indicates up-regulated DEGs, while the lower row represents down-regulated DEGs. Red numbers indicate DEGs exclusively shared in either the resistant or susceptible lines. (c) Overrepresented GO terms of 204 up-regulated DEGs exclusively in the resistant lines at 8 hpi. (d) Overrepresented GO terms of 308 up-regulated and 42 down-regulated DEGs exclusively in the resistant lines at 16 hpi. (e) Fold changes in H_2_O_2_ concentration after *P. sansomeana* inoculation at 16 hpi. Log_2_ fold change was calculated based on H_2_O_2_ concentration between mock- and pathogen-inoculated plants at 16 hpi. GO terms analysis (refer to Figure 2d) identified the enrichment of H_2_O_2_ catabolic process and reactive oxygen species metabolic process for up-regulated genes in the resistant lines (Colfax and NE2701) after pathogen inoculation. The error bars represent SD (standard deviation) among three separate measurements. (f) Overrepresented GO terms of 22 up-regulated and 4 down-regulated DEGs exclusively in the susceptible lines at 4 hpi. For panels c, d, and f, the numbers within the bars denote the count of genes enriched in each GO term (BP, Biological process; CC, Cellular component; MF, Molecular function; KEGG, Kyoto encyclopedia of genes and genomes).

To specifically identify DEGs unique to either the resistant or susceptible lines, we performed an intersection analysis of these DEGs between different soybean lines (Figure 2b). In the early stages at 2 and 4 hpi, only a limited number of DEGs were common in the two resistant lines. However, at 8 and 16 hpi, a substantial overlap was observed, with several hundred up-regulated DEGs being shared between Colfax and NE2701, while only a few dozen down-regulated DEGs were common to these two lines. Among the DEGs identified in the resistant lines, we identified *Glyma.09G210600* (Dataset S1), a homolog of *resistance to pseudomonas syringae 3* (*RPS3*) in Arabidopsis, which encodes a NBS-LRR-type protein, known for its role as a R protein in the defense response (Dangl & Jones, 2001; Lin et al., 2014; Sandhu et al., 2005; van der Hoorn & Kamoun, 2008). Interestingly, it showed significant up-regulation in response to the pathogen inoculation at 8 or 16 hpi exclusively in both resistant soybean lines (Figure S3; Dataset S1).

To elucidate the function of these DEGs, we performed gene ontology (GO) enrichment analysis. Enriched GO terms were only identified at 8 and 16 hpi in the resistant lines, and at 4 hpi in the susceptible lines. At 8 hpi, the 204 up-regulated DEGs uniquely shared by the two resistant lines were primarily associated with ET-responsive functions, including the ethylene-activated signaling pathway (GO:0009873), cellular response to ethylene stimulus (GO:0071369), and response to ethylene (GO:0009723) (Figure 2c). In the molecular function category, we identified 24 genes exhibiting DNA-binding transcription factor activity (GO:0003700) that were exclusively up-regulated in the resistant lines at 8 hpi. These include multiple ethylene response factors (*ERF*s) and various other transcription factors. Moreover, in the Kyoto encyclopedia of genes and genomes (KEGG), the MAPK signaling pathway (KEGG:04016) was significantly overrepresented among the up-regulated genes at 8 hpi in the resistant lines.

At 16 hpi, the 308 commonly up-regulated DEGs in the resistant lines demonstrated the most significant enrichment in GO terms related to defense response (GO:0006952), response to stress (GO:0006950) or stimulus (GO:0050896), along with several other terms associated with stress or defense responses, such as hydrogen peroxide catabolic process (GO:0042744) and ROS metabolic process (GO:0072593) (Figure 2d). To illustrate the accumulation of hydrogen peroxide (H_2_O_2_) accumulated in the resistant lines, we quantified the concentration of H_2_O_2_ in the four lines at 16 hpi. We observed an increase in the concentration of H_2_O_2_ in the two resistant lines upon *P. sansomeana* inoculation at 16 hpi (Figure 2e). In contrast, the two susceptible lines exhibited a lower concentration of H_2_O_2_ under pathogen inoculation compared to mock inoculation.

In contrast, the susceptible lines had fewer common DEGs across all time points. Notably overrepresented GO terms in the susceptible lines were only detected at 4 hpi (Figure 2f). The majority of the significantly enriched GO classes for the down-regulated DEGs were predominantly associated with photosynthesis-related classes, including photosystem I (GO:0009522) and photosystem II (GO:0009523), and photosynthesis light harvesting (GO:0009765).

In addition to the differential transcriptomic responses between the resistant and susceptible lines, we also observed distinctive defense responses within the two resistant lines. In NE2701, the highest number of genes exhibited differential regulation as early as 2 hpi, in contrast to Colfax, where the most significant differences in DEGs emerged at a relatively later time point, 16 hpi (Figure 2a,b). Enriched GO terms in the biological process category for DEGs exclusively at 2 hpi in NE2701 included regulation of the jasmonic acid mediated signaling pathway (GO:2000022) and plant-type cell wall organization (GO:0009664) (Table S1). On the other hand, in Colfax, up-regulated genes were most significantly enriched for the phosphate-containing compound metabolic process (GO:0006796), while down-regulated genes were notably associated with photosynthesis (GO:0015979) at 16 hpi. These findings suggest that in addition to common defense strategies shared by these two resistant lines, genetic variations between these two lines also contribute to the distinct transcriptomic changes in response to the pathogen *P. sansomeana*.

### Ethylene-responsive genes are mainly co-expressed in the two resistant lines in response to the pathogen inoculation

To identify co-expressed genes in response to the pathogen across the four lines, we extracted gene clusters demonstrating similar patterns of gene regulation under different conditions for each line. From these grouped clusters in each line (Figure S4), we selected the clusters likely to contain co-expressed genes that were either up-regulated or down-regulated upon pathogen inoculation (Figure 3; Dataset S2). However, no clusters from Senaki and Williams 82 displayed down-regulation in response to the pathogen inoculation. Therefore, we focused only on the clusters associated with up-regulation (Figure S4). We identified two clusters (503 and 1606 genes; Figure 3a) in Colfax, along with one cluster for each of the other soybean lines, NE2701 (1091 genes; Figure 3b), Senaki (1891 genes; Figure 3c), and Williams 82 (981 genes; Figure 3d), all exhibiting up-regulation upon pathogen inoculation.

**Figure 3.**
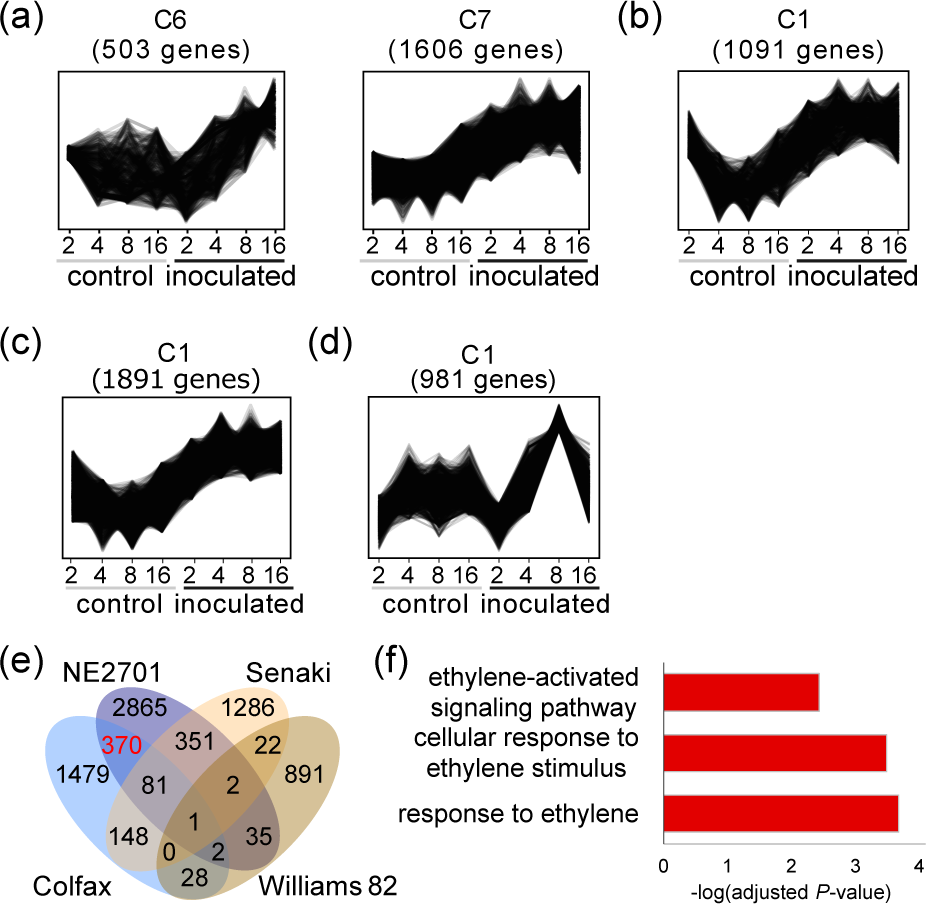
Co-expressed gene clusters up-regulated upon pathogen inoculation in the resistant lines are enriched in ethylene-responsive genes. (a-d) Co-expression gene clusters demonstrating similar patterns of up-regulation upon pathogen inoculation in Colfax (a), NE2701 (b), Senaki (c), and Williams 82 (d). These clusters were selected from the overall clusters, grouped based on *k*-means clustering (see Figure S4). The X-axis indicates time points: 2, 4, 8, and 16 hpi. The detailed gene list for each selected cluster can be found in Dataset S2, with enriched GO terms available in Dataset S3. (e) Shared and unique up-regulated DEGs within clusters across the four lines after inoculation. (f) Overrepresented GO terms in the gene lists of shared genes exclusively in the resistant lines. A total of 370 genes marked in red in (e) were used for the GO analysis. All three GO terms are classified under the biological process category.

In these gene clusters, numerous GO classes associated with diverse biological processes and molecular functions were significantly enriched (Dataset S3). To pinpoint co-expressed genes specific to the resistant lines, we intersected the genes from the selected up-regulated clusters and extracted 370 genes exclusively present in the resistant lines (Figure 3e). Among these genes, ET-responsive genes (response to ethylene, GO:0009723; cellular response to ethylene stimulus, GO:0071369; ethylene-activated signaling pathway, GO:0009873) were overrepresented as the main globally co-expressed genes up-regulated in response to the pathogen inoculation in the resistant lines (Figure 3f). These genes included several ethylene response factors (ERFs) like *ERF1*, *ERF15*, and *ERF98*, known for their regulatory role in ET-responsive genes (Gao et al., 2020; Thirugnanasambantham et al., 2015). Additionally, the ethylene-forming enzyme *1-aminocyclopropane-1-carboxylate oxidase 4* (*ACO4*) was part of this ET-responsive gene network (Table S2).

### The majority of the differentially transcribed TEs are up-regulated in response to *P. sansomeana* in all lines, and those in Colfax are more enriched near genes

As TEs are often subject to environmental changes (Casacuberta & González, 2013), we compared transcriptional patterns of TEs in the four soybean lines in response to *P. sansomeana*. Based on the annotation of the soybean reference genome (Wm82.a4), we examined the expression of TEs across ten superfamilies, including four retrotransposons (class I) and six DNA transposons (class II) (Figure 4; Figure S5). Out of the 324,333 TEs, only an average of 3% exhibited transcription, in either mock-inoculated or pathogen-inoculated plants. Notably, although long interspersed nuclear elements (LINEs) were not the most abundant TE superfamily in the soybean genome, they constituted the largest proportion of transcribed TEs at 15% compared to other TE superfamilies (Figure S5). In Colfax, more TEs in most superfamilies were transcribed upon the pathogen inoculation compared to the mock inoculation consistently at 2 hpi, and predominantly at 16 hpi, particularly in DNA transposons. In Senaki, a similar overall increase in transcribed TEs upon inoculation was observed at 2 hpi, mirroring the trend noted in Colfax, except for DNA/*TcMar*/*Stowaway*, which exhibited reduced activation upon inoculation (Figure S5e). Overall, no significant differences in the total transcript levels of TEs were observed between the resistant and susceptible lines across most superfamilies (Figure S6). Interestingly, DNA/*PIF*-*Harbinger* showed higher transcript levels in both resistant lines at 2 hpi post the *P. sansomeana* inoculation, while the susceptible lines displayed comparable transcript levels between mock-inoculated and pathogen-inoculated plants (Figure S6d).

**Figure 4.**
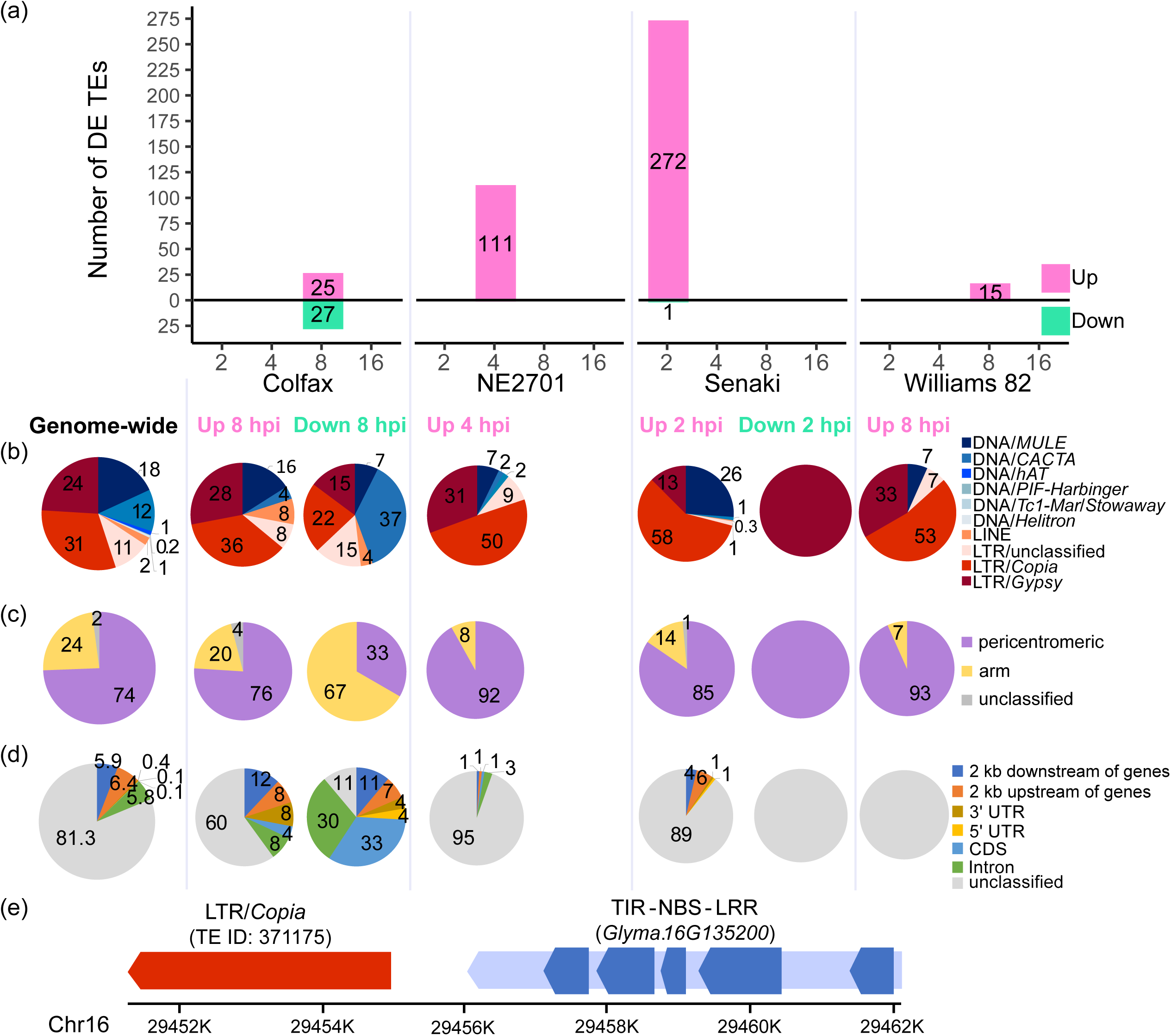
The majority of differentially expressed TEs are up-regulated in response to *P. sansomeana* and those in Colfax are enriched near genes. (a) The number of up- and down-regulated TEs in four lines upon inoculation at different time points (2, 4, 8, and 16 hpi). (b) The percentage of differentially expressed (DE) TEs in each line. The term “genome-wide” indicates the TE composition in the soybean reference genome. Retrotransposons (class I) are represented by four families in reddish colors, while DNA transposons (class II) are represented by six superfamilies in bluish colors. (c) The percentage of DE TEs between pericentromeric and chromosomal arms. The leftmost pie indicates the locations of all TEs in the genome, serving as a control. (d) The percentage of DE TEs near or within genes. Each pie, except the “genome-wide” in (b-d), corresponds to the respective soybean line in (a). The numbers in (b-d) denote the respective percentages. (e) Diagram illustrating the position of a TE (LTR/*Copia*) downstream of the *TNL* gene on chromosome 16. Both the TE and gene were up-regulated in Colfax upon pathogen inoculation. The dark blue boxes indicate the five exons in the *TNL* gene.

Next, we identified significantly differentially expressed TEs (DE TEs) between the mock and pathogen inoculated samples at different time points for each of the soybean lines (Dataset S4). The number of DE TEs varied among the four lines in response to the pathogen inoculation (Figure 4a). In Colfax, comparable numbers of up-regulated and down-regulated DE TEs (25 and 27, respectively) in response to the pathogen were detected only at 8 hpi. The proportion of TE superfamilies among the up-regulated TEs in Colfax mirrored the genome-wide level in the soybean genome, where approximately 70% of TEs represented retrotransposons and the remaining 30% were DNA transposons (Figure 4b). These up-regulated TEs were located mainly in pericentromeric regions (76%), similar to the genome-wide distribution (74%) (Figure 4c). Furthermore, more than 40% of the up-regulated DE TEs in Colfax were located proximal to genes in the genome (Figure 4d). In contrast, a larger proportion of the down-regulated DE TEs in Colfax consisted of DNA transposons (44%), particularly *CACTA* elements (37%). Interestingly, over two-thirds of the down-regulated DE TEs in Colfax were located in chromosomal arms, and a significant portion (90%) of these down-regulated DE TEs in Colfax were either within or in close proximity to genes (within 2 kb upstream or downstream), a marked contrast to the overall genome-wide distribution where less than 20% of TEs were positioned near genes. In Colfax, among the genes near or containing up-regulated DE TEs, a significantly up-regulated gene (*Glyma.16G135200*) exhibited an association with an up-regulated LTR/*Copia* element (ID 371175) (Figure 4e; Dataset S1). This gene encodes a Toll/interleukin-1-receptors (TIR)-NBS-LRR (TNL) protein (Swiderski et al., 2009), renowned for its regulatory role as a R protein in immune responses to *Phytophthora sojae* in soybean (Zhou et al., 2022). Additionally, in Colfax, among the DEGs with down-regulated DE TEs, three harbored DNA/*CACTA* elements within their coding regions (Dataset S4). Notably, two out of these three DEGs (*Glyma.19G016400* and *Glyma.13G063700*) were both ABC transporters that were also down-regulated upon inoculation (Dataset S1).

In NE2701, a considerably larger number of DE TEs (n=111) were up-regulated at a relatively earlier stage, 4 hpi (Figure 4a), compared to Colfax. The majority of up-regulated DE TEs in NE2701 were retrotransposons (90%), including LTR/*Copia* accounting for approximately half of all TEs (Figure 4b). Most (92%) of the up-regulated DE TEs in NE2701 were located in pericentromeric regions, a proportion much higher than the overall genome-wide distribution (74%) (Figure 4c). Furthermore, in contrast to Colfax, the up-regulated DE TEs in NE2701 (95%) were primarily located outside gene regions (Figure 4d).

In the susceptible line, Senaki, a total of 272 DE TEs exhibited up-regulation at 2 hpi, while only one TE displayed down-regulation at the same time point (Figure 4a). Over half of the up-regulated DE TEs (58%) in Senaki were LTR/*Copia*, unlike the other lines and the genome-wide proportion. Nonetheless, the overall proportions of classes I and class II TEs were comparable to the genome-wide levels (Figure 4b). In the other susceptible line, Williams 82, a lower number of DE TEs (15) were observed to be up-regulated at 8 hpi. Out of the 15 up-regulated DE TEs, 14 were retrotransposons. The DE TEs in both susceptible lines were primarily located in pericentromeric regions (85% and 93%, respectively) and outside gene regions (89% and 100%, respectively) (Figure 4c,d). Taken together, these data propose that pathogen attacks trigger the transcriptional activation of numerous TEs, and the transcriptional responses of TEs to *P. sansomeana* vary significantly across different soybean lines.

### LncRNA XLOC_013220 is associated with the expression of its potential target genes following pathogen inoculation in the resistant lines

LncRNAs have been reported to regulate gene expression in various biological processes including pathogen infection (Chekanova, 2015; Gil & Ulitsky, 2020). To investigate how soybean lncRNAs respond to *P. sansomeana* and their potential interaction with gene expression, we first identified lncRNAs in soybean using the assembled and merged transcripts from the 64 samples. Our analysis revealed a total of 43,759 lncRNA transcripts, comprising long intergenic non-coding RNAs (lincRNAs), sense-overlapping lncRNA transcripts (SOT), and antisense-overlapping lncRNA transcripts (AOT) (Table S3). The vast majority of the identified lncRNAs were lincRNAs (95%; 41,159), with a smaller proportion representing SOT or AOT (3% and 2%, respectively) (Figure 5a). Additionally, based on sequence similarities with TEs, we classified lincRNAs into non-TE lincRNAs and TE-containing lincRNAs, possibly indicating historical TE integration or exaptation events (Nelson et al., 2017). Among the identified lncRNAs, approximately 58% (27,672) were designated as TE-containing lincRNAs, while the remaining 37% (13,487) were classified as non-TE lincRNAs (Figure 5a).

**Figure 5.**
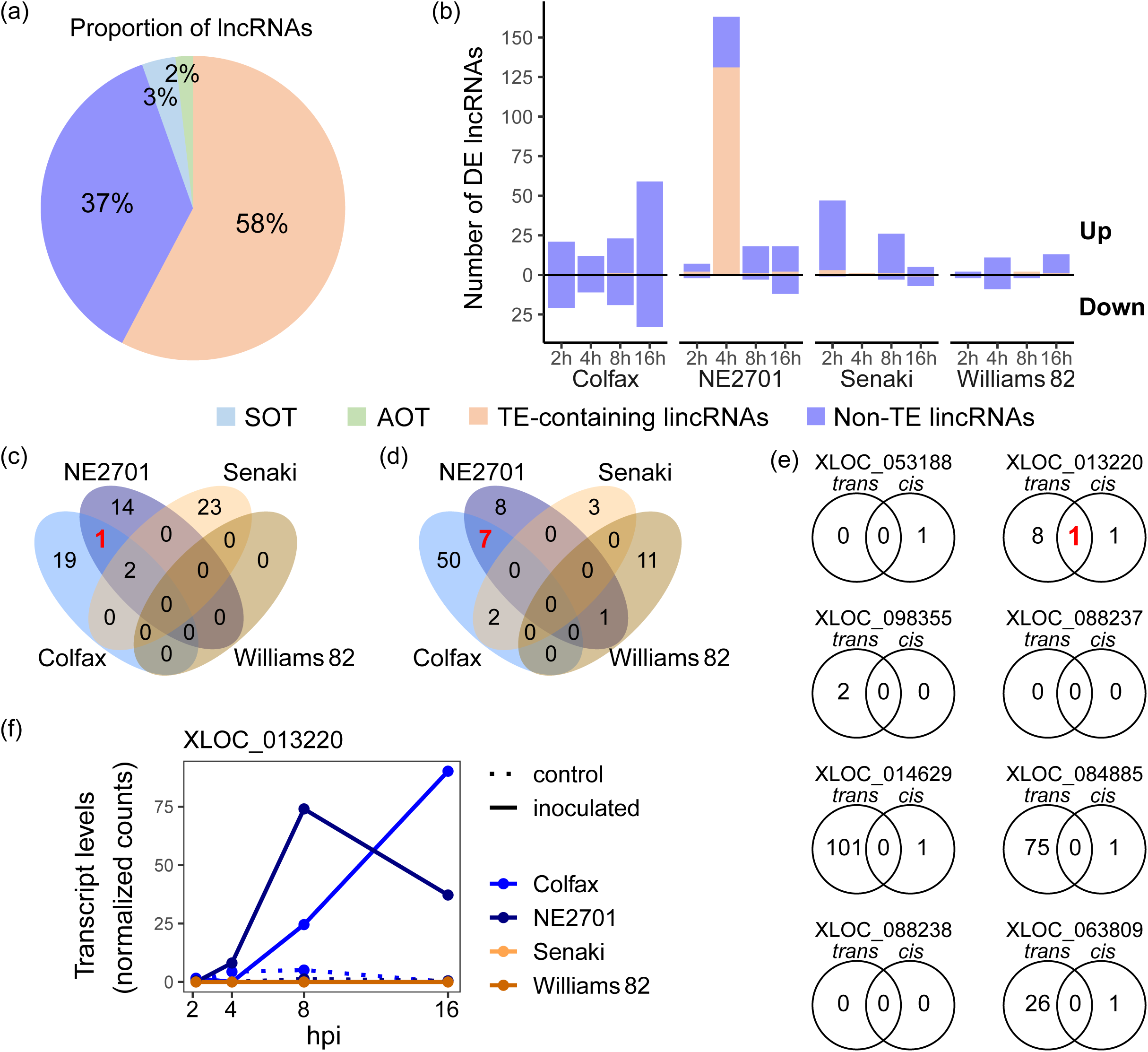
LncRNA XLOC_013220 is associated with the expression of its potential target genes following pathogen inoculation in the resistant lines. (a) The proportion of identified lncRNAs in the soybean transcriptomes. (b) The number of up- and down-regulated lincRNAs upon inoculation in each line at different time points (*P* < 0.01). (c-d) The number of up-regulated non-TE lincRNAs at 8 hpi (c) and 16 hpi (d), where shared lincRNAs were exclusively found in either the resistant or susceptible lines. The shared lincRNAs were only present in the resistant lines as up-regulated at 8 and 16 hpi (no shared down-regulated lincRNAs were identified). The red number represents the shared DE non-TE lincRNAs in the resistant lines. (e) The number of potential *trans*-and *cis*-target genes for each lincRNAs identified as shared DE non-TE lincRNAs in the resistant lines in (c) and (d). The red number indicates the gene *Glyma.03G040900*, which encodes a LURP-one-related (LOR) protein and is recognized as both the potential *trans*-and *cis*-target of XLOC_013220. (f) Transcript level change of XLOC_013220 at different time points for each line and condition.

We next compared the proportion of transcription and transcript levels among non-TE lincRNAs, TE-containing lincRNAs, genes, and TEs. Among these four transcript types, transcribed genes showed the highest proportion, accounting for 74% of the total transcripts (Figure S7a). In the context of lincRNAs, non-TE lincRNAs exhibited a higher transcription proportion at 15%, in contrast to TE-containing lincRNAs with 12%. In contrast, TEs demonstrated the lowest transcription proportion at 3% compared to the other four transcript types. This trend was consistent when considering mean transcript levels, where transcribed genes exhibited significantly higher levels compared to the other transcript types (Figure S7b). Similarly, within the other three transcript types, non-TE lincRNAs exhibited higher mean transcript levels, while TEs displayed the lowest mean transcript levels, characterized by relatively larger variability compared to lincRNAs (Figure S7b).

Upon the *P. sansomeana* inoculation, we observed differential expression in a total of 497 unique lincRNAs compared to mock-inoculated plants (Figure 5b; Dataset S5). The majority of differentially expressed lincRNAs (96%) were non-TE lincRNAs across all four soybean lines, except for NE2701 at 4 hpi, where 131 TE-containing lincRNAs were up-regulated. Of these 131 TE-containing lincRNAs, 38 (29%) directly overlapped with DE TEs. In contrast, both susceptible lines, Senaki and Williams 82, showed fewer numbers of DE lincRNAs, most of which were up-regulated following inoculation (Figure 5b).

We further investigated common lincRNAs exclusively present in either the resistant or susceptible lines. Remarkably, we identified eight non-TE lincRNAs exclusively up-regulated in the resistant lines at 8 and 16 hpi (Figure 5c,d). A correlation analysis between these eight lincRNAs and DEGs revealed a wide range of co-expressed genes with each lincRNA (Figure 5e; Dataset S6). Among these, XLOC_013220 and XLOC_098355 were linked to nine and two co-expressed DEGs, respectively. Additional non-TE lincRNAs, such as XLOC_014629, XLOC_084885, and XLOC_063809, exhibited a relatively higher number of co-expressed DEGs. The enriched GO terms for the DEGs associated with XLOC_014629 and XLOC_084885 primarily pertained to the biosynthesis of various secondary metabolites. In contrast, DEGs involved in salicylic acid biosynthesis were enriched for XLOC_063809 (Table S4).

We also explored potential *cis*-target genes by identifying adjacent genes (± 5 kb) to the eight non-TE lincRNAs in the resistant lines (Figure 5c,d). Among these lincRNAs, four lincRNAs (XLOC_053188, XLOC_014629, XLOC_084885, and XLOC_063809) had a single adjacent gene each (Dataset S7). However, these genes were not DEGs (Dataset S1). Remarkably, an intriguing lincRNA, XLOC_013220, located between two flanking genes (*Glyma.03G040900* and *Glyma.03G041000*), both encoding LURP-one-related (LOR) proteins, exhibited a significant correlation with the transcript level of these two genes (Figure 5e). LORs have been demonstrated to play a role in defense responses to oomycete pathogens in Arabidopsis (Baig, 2018; Knoth & Eulgem, 2008). Interestingly, both of these *LOR* genes were DEGs upon pathogen inoculation at both 8 and 16 hpi exclusively in the resistant lines (Dataset S1). Significantly, *Glyma.03G040900* was also identified as a potential *trans*-target gene, showing a strong correlation in transcript levels with the lincRNA (R = 0.85, *P* = 2.2E-16). The up-regulation of this gene after pathogen inoculation in the resistant lines was further validated by qRT-PCR (Figure S8). Similarly, the transcript levels of the other *LOR* gene, *Glyma.03G041000*, also demonstrated a significant correlation with the lincRNA (R = 0.63, *P* = 2.2E-08). This particular lincRNA, XLOC_013220, exhibited exclusive up-regulation in the resistant lines at 16 hpi, with a consistent trend also at 8 hpi, aligning with the expression patterns of the two flanking genes (Figure 5f; Dataset S1). This lincRNA is of particular interest as it is located within the mapped region of the resistance gene in the resistant line Colfax based on our mapping results (Lin et al., 2023). These findings strongly suggest that XLOC_013220 potentially plays a crucial role in regulating the expression of both neighboring *LOR* genes in response to the pathogen infection in the resistant lines.

Among the DE TE-containing lincRNAs, only one lincRNA was exclusively shared by the resistant lines (Figure S9). This lincRNA harbored a DE *CACTA* element (ID 294229, Dataset S4) embedded directly within its transcript region. Furthermore, we detected 49 DEGs that exhibited strong co-expression patterns with this specific lincRNA based on the correlation analysis (Dataset S8).

### CHH methylation levels in lincRNAs increase upon pathogen inoculation at the later time point in the resistant line compared to the susceptible line

LncRNAs have been found to interact with DNA methyltransferase to meditate gene expression (Böhmdorfer et al., 2014; Zhao et al., 2016). In pursuit of a more profound understanding of the prospective role of lincRNAs in gene expression modulation, we investigated DNA methylation levels within non-TE lincRNAs between the resistant line Colfax and the susceptible line Williams 82. We first determined DNA methylation levels in the CG, CHG and CHH contexts within lincRNA transcript bodies, as well as in regions extending 2 kb upstream of the start positions and 2 kb downstream of the end positions. Strikingly, across all regions and in each of the three cytosine contexts, the resistant line Colfax demonstrated marginally lower methylation levels in lincRNAs compared to the susceptible line Williams 82 (Figure 6 and Figure S10), suggesting distinct methylation patterns in lincRNAs across different soybean genetic backgrounds.

**Figure 6.**
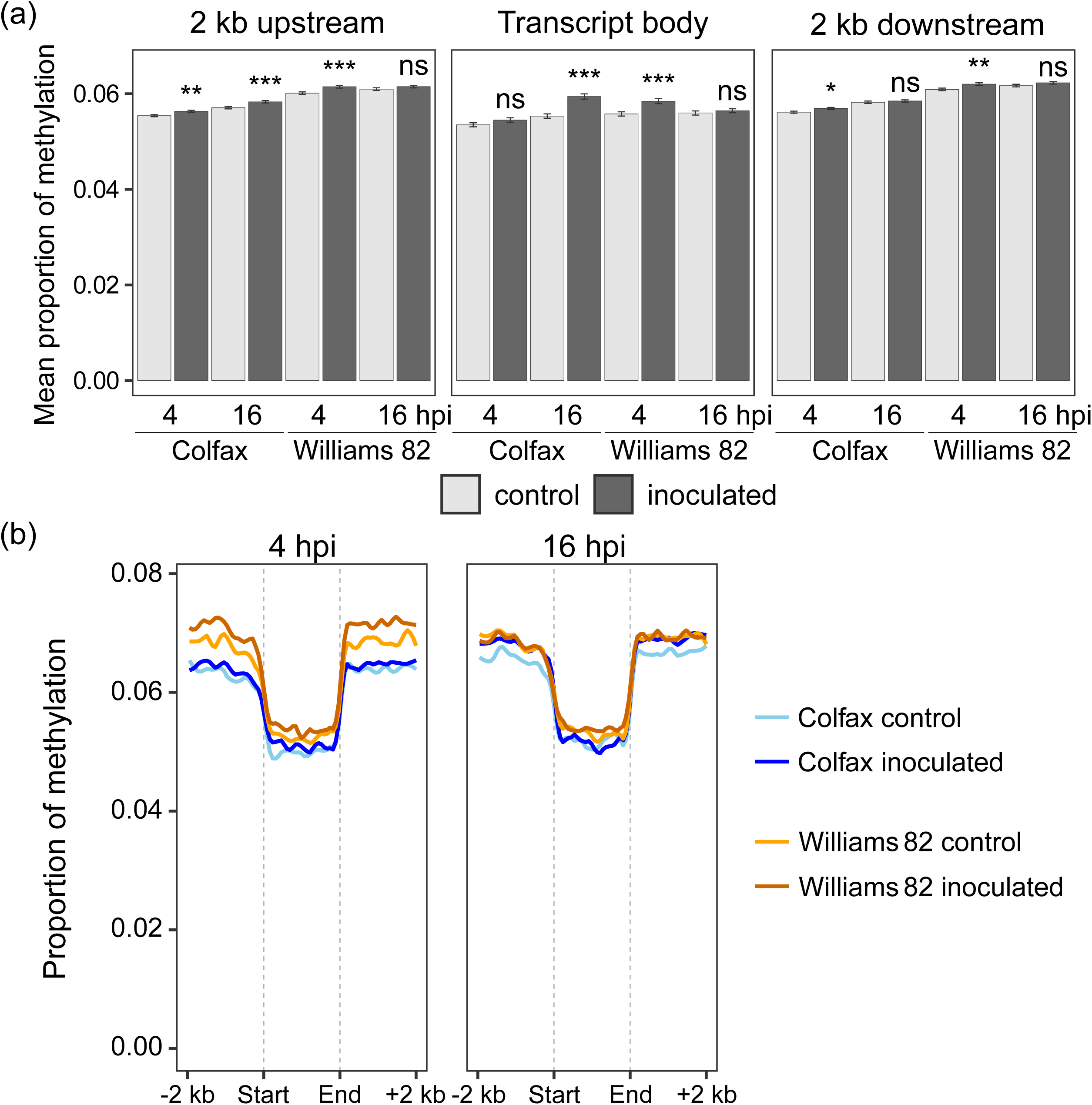
CHH methylation levels in lincRNAs increase upon pathogen inoculation at the later time point in the resistant line compared to the susceptible line. (a) Mean proportions of CHH methylation within non-TE lincRNA transcript bodies, and 2 kb up or downstream of these bodies for each line and condition. The error bars depict ± SE. Asterisks denote statistically significant differences in means between inoculated plants and controls (Student’s *t* test; ***, *P*Lvalue < 0.001; **, *P* < 0.01; *, *P* < 0.05; ns, not significant, *P* ≥ 0.05). (b) Distribution of CHH methylation within 2 kb upstream and downstream of non-TE lincRNA transcript bodies. The mean methylation proportion was calculated in 40 windows for each of upstream, body, and downstream of transcripts. The lines were smoothed using locally estimated scatterplot smoothing (LOESS).

To further understand the role of methylation in response to the pathogen, we examined the methylation changes upon pathogen inoculation. While no significant methylation changes in CG and CHG contexts were observed after pathogen inoculation in either the resistant or susceptible lines within 2 kb flanking regions and lincRNA bodies (Figure S10), CHH methylation levels increased in both lines following inoculation (Figure 6). Interestingly, the CHH methylation pattern and timing of response to the pathogen inoculation varied between the two lines. In the resistant line Colfax, CHH methylation levels in lincRNAs exhibited a more pronounced increase at 16 hpi, particularly within the transcript bodies (Figure 6a). In contrast, in the susceptible line Williams 82, CHH methylation levels in lincRNAs showed a more pronounced increase at the earlier time point of 4 hpi. The distribution of CHH methylation across 2 kb flanking regions also showed earlier increases at 4 hpi in the susceptible line and delayed changes at 16 hpi in the resistant line (Figure 6b). These results suggest that lincRNAs in the resistant line may possess a relatively higher stability with respect to epigenetic changes in response to pathogen attack compared to those in the susceptible line.

## Discussion

### *P. sansomeana* is likely a hemibiotrophic oomycete pathogen

Our transcriptomic analyses demonstrated that genes involved in the ET signaling pathway were significantly up-regulated during later stages of *P. sansomeana* inoculation (8 and 16 hpi) in the resistant lines (Figures 2, 3, and Table S2). This coordinated activation likely interacts with the up-regulation of multiple transcription factors that initiate and modulate diverse defense responses. Among these transcription factors, *ERF*s have well-documented roles in intricate networks involving ET and other hormones to enhance stress tolerance in plants (Adie et al., 2007; Dubois et al., 2018; Thirugnanasambantham et al., 2015). Several up-regulated *ERF* genes associated with the ET signaling exhibited a co-expression pattern in response to *P. sansomeana* (Figure 3).

ET, in concert with JA, is a key signal molecule in host defense response against necrotrophic and later-stage hemibiotrophic pathogens (Huang et al., 2020; van Loon et al., 2006). Clear evidence shows that many *Phytophthora* spp. pathogens, such as *Phytophthora infestans* and *P. sojae*, follow a hemibiotrophic life cycle, initiating the infection cycle as biotrophs but switching to a necrotrophic lifestyle at later stages (Lee & Rose, 2010; Qutob et al., 2002; Zuluaga et al., 2016). In *P. sojae*, an avirulence (effector) gene product with an RxLR (Arginine-any amino acid-Leucine-Arginine) motif can interact with a corresponding soybean *Rps* gene, resulting in the rapid host activation of defense responses and plant resistance (Arsenault-Labrecque et al., 2022; Dong et al., 2011; J. D. Jones & J. L. Dangl, 2006; Na et al., 2013; Ngou et al., 2022; Shan et al., 2004). As disease progresses, *P. sojae* feeds on dead plant tissues, causing severe lesions and leading to necrosis (Qutob et al., 2002). According to our mapping results, a single dominant resistance gene contributes major resistance to *P. sansomeana* (Lin et al., 2023), suggesting that *P. sansomeana* possesses similar genetic qualities, such as effectors. The up-regulation of the ET signaling pathway and the ROS metabolic process at 8 and 16 hpi from our transcriptomic data and the H_2_O_2_ experiment demonstrated necrotrophic symptoms. Overall, these findings suggest that, similar to other *Phytophthora* spp. pathogens, *P. sansomeana* is likely a hemibiotrophic pathogen. Future investigation into the life cycle of *P. sansomeana* could provide additional evidence.

### LincRNA XLOC_013220 potentially regulates adjacent *LOR*s upon pathogen invasion

LncRNAs can regulate diverse defense mechanisms by influencing the expression of resistance-related genes at both transcriptional and post-transcriptional levels (Sharma et al., 2022). In this study, we identified several hundreds of *trans*-targets for DE lincRNAs upon pathogen inoculation in the resistant lines, providing potential candidate target genes or co-expressed downstream genes that are induced in response to *P. sansomeana* (Figure 5). These lincRNAs may directly control or indirectly influence pathways involved in defense responses, such as lignan and SA biosynthesis (Table S4).

Of these DE lincRNAs that co-expressed with DEGs, an especially intriguing instance involves the lincRNA XLOC_013220, which experienced up-regulation upon pathogen inoculation. This lincRNA is of particular interest as it lies within the mapped region of the resistance gene in the resistant line Colfax (Lin et al., 2023). More interestingly, it is positioned adjacent to the two *LOR* genes, presumably generated by tandem duplication. *LURP1*, previously identified to be up-regulated upon recognition of pathogenic oomycetes in plants, is a key gene controlling basal defense against the oomycete species *Hyaloperonospora parasitica* in *Arabidopsis thaliana* (Knoth & Eulgem, 2008). It is worth noting that *LUPR1* in *A. thaliana* possesses a W-box and two TGA-box motifs that might engage with members of the WRKY family, which play roles in both PTI and ETI responses. In addition, a *LOR* gene, belonging to the *LURP* cluster, has been recently unveiled as a contributor to basal defense against another oomycete, *Hyaloperonospora arabidopsis* (Baig, 2018). In Arabidopsis, both *LURP1* and *LOR* are involved in the SA-dependent pathway, orchestrating immune responses to these oomycete pathogens.

Given the close proximity and the highly correlated expression patterns between the lincRNA and the two *LOR* genes in the resistant lines, it is possible that this lincRNA can exert a regulatory role in the expression of its neighboring *LOR* genes, thereby contributing to defense responses in the resistant soybean lines against the oomycete pathogen *P. sansomeana*. The lincRNA could function as a *cis*-regulatory element, modulating the transcriptional activity or stability of the *LOR* genes (Gil & Ulitsky, 2020). Alternatively, the lincRNA might be involved in coordinating the expression of these genes within a larger regulatory network activated in response to the pathogen. Further investigation, such as functional studies or genetic manipulations, would be necessary to elucidate the precise role of the lincRNA in regulating the expression of the neighboring *LOR* genes during the pathogenic response.

### Resistant and susceptible lines exhibit distinct patterns of CHH methylation in lincRNAs against *P. sansomeana*

Our methylation data revealed that CHH methylation levels in lincRNAs and their flanking regions in response to the *P. sansomeana* inoculation increased at distinct time points in both the resistant and susceptible lines, while the CG and CHG methylation levels remained unchanged (Figure 6; Figure S10). Although the methylation levels of CHH cytosines are generally very low, they are remarkably abundant in the soybean genome (Song et al., 2013), making them potential candidates for buffering the global impact of environmental stresses, such as pathogen attacks, on transcriptional activation of TEs to maintain genome stability. Interestingly, although both the resistant and susceptible lines exhibited increased CHH methylation levels in response to *P. sansomeana*, the timing of this increase differed between them (Figure 6). In the resistant line, a more significant increase in CHH methylation levels in lincRNAs occurred at 16 hpi. In contrast, the susceptible line exhibited this increase at an earlier time point, 4 hpi. This divergence in timing may be attributed to their distinct genetic backgrounds or variations in the kinetics of their defense response. The resistant line seems to mount a delayed yet sustained CHH methylation response, contributing to a more gradual and prolonged defense response. On the other hand, the susceptible line experienced an earlier but potentially transient increase in global CHH methylation triggered by *P. sansomeana* infection.

### Distinct genetic backgrounds may contribute to divergent defense strategies between the two soybean resistant lines against *P. sansomeana*

In addition to common defense responses shared by both resistant lines, we further dissected unique strategies employed by each resistant line. In the resistant line Colfax, notable transcriptomic changes, spanning genes, TEs, and lincRNAs, primarily occurred at 8 or 16 hpi in response to the pathogen. At 16 hpi, an exclusive and significantly enriched biological process in Colfax for the up-regulated genes was the phosphate-containing compound metabolic process, which encompasses crucial phosphorylation events (Table S1). This process might play a role in activating *ERF*s, crucial in ET-mediated immune responses (Adie et al., 2007; Thirugnanasambantham et al., 2015; Wang et al., 2022). By contrast, the resistant line NE2701 exhibited larger proportions of differentially transcribed genes, TEs, and lincRNAs at relatively earlier time points, 2 or 4 hpi. Notably, the primary biological process initiated immediately after pathogen attack at 2 hpi in NE2701 was the JA-mediated defense signaling pathway being triggered.

The divergent patterns of defense responses between the two resistant lines were also evident in differentially transcribed TEs and lncRNAs. In Colfax, DE TEs were largely located within gene regions (Figure 4d). Of particular interest was the up-regulated TE (LTR/*Copia*) located closely downstream of the significantly up-regulated *TNL* gene (*Glyma.16G135200*), which showed a similar gene expression pattern to the TE (Figure 4e; Dataset S1). Recently, a *TNL* was newly identified in soybean, demonstrating resistance against Phytophthora root rot (Zhou et al., 2022), and has been shown to increase JA- and SA-mediated disease resistance. Our finding suggests a potential role for this TE as a *cis*-regulatory element, modulating the expression of the *TNL* gene in defense responses to the pathogen in Colfax.

On the other hand, the NE2701 resistant line showed a higher number of up-regulated TEs and lincRNAs in response to *P. sansomeana*, particularly at the relatively earlier time point, 4 hpi (Figures 4 and 5). This suggests a more rapid alleviation of silencing of TEs in NE2701 in response to the pathogen compared to Colfax. Strikingly, unlike Colfax, these DE TEs in NE2701 were located predominantly outside gene regions. Interestingly, over one-third of the up-regulated TEs in NE2701 overlapped with lincRNAs, which were also up-regulated post inoculation, suggesting that some of these TEs could serve as sources for lincRNAs that modulate transcriptomic responses to the pathogen (Sharma et al., 2022; Zhang et al., 2020).

It is worth noting that the different timing and patterns of transcriptional responses between the two resistant lines can be attributed to their unique genetic backgrounds. Given that NE2701 is originally derived from a cross between Colfax and A91-701035, it is most likely that they share the same *R* gene responsible for the resistance against *P. sansomeana*. However, A91-701035 has a divergent genetic background due to consecutive crosses with multiple other soybean lines (Graef et al., 2005). As a result, there may be some additional minor alleles from A91-701035 that also contribute to the disease response, in addition to the *R* gene inherited from Colfax. These unique genetic backgrounds likely shape the defense strategies observed in each of the resistant lines, highlighting the potential benefits of leveraging diverse genetic backgrounds for breeding more resilient and effective soybean cultivars against pathogenic challenges.

## Acknowledgements

We thank HiPerGator Supercomputer for providing us with the computational resources to perform the analysis.

## Funding

This work was supported by the National Science Foundation under Award Number IOS2128023, the National Institute of General Medical Sciences of the National Institutes of Health under Award Number R15GM135874, as well as startup funds from Miami University and the University of Florida to M.Z. We also express our gratitude for partial support from the Michigan Soybean committee, North Central Soybean Research Program, and Project GREEEN-Michigan’s plant agriculture initiative to M.I.C. Additionally, we thank funding support from the Michigan Soybean Committee, Project GREEEN-Michigan’s plant agriculture initiative, AgBioResearch at Michigan State University (Project No. MICL02013), North Central Soybean Research Program, United States Department of Agriculture National Institute of Food and Agriculture (Hatch project 1011788), and the United Soybean Board (24-209-S-A-1-A) to D.W. The funders had no role in study design, data collection and analysis, decision to publish, or preparation of the manuscript.

**Competing interests:** The authors declare no conflict of interest.

## Data Availability Statement

The raw and processed data of mRNA and whole genome bisulfite sequencing presented in this study have been deposited in NCBI Gene Expression Omnibus under the accession number GSE240966, https://www.ncbi.nlm.nih.gov/geo/query/acc.cgi?acc=GSE240966.

## Supporting information

**Figure S1. Phenotypes of mock- and *P. sansomeana*-inoculated plants of the four soybean lines.** The photos were taken five days after mock- or pathogen-inoculation for four lines.

**Figure S2. Principal component analysis (PCA) of 64 soybean transcriptomes.** Variant stabilizing transformations of read counts were used for PCA to determine the sample distance in four soybean lines. The percentages of variation explained by PC1 and PC2 are indicated in parentheses.

**Figure S3. Transcript levels of *Glyma.09G210600* (*RPS3*), exclusively up-regulated at 16 hpi in both resistant lines.** (a) Transcript levels of *RPS3* from RNA sequencing. Normalized counts were used for transcript levels at different time points for each line and condition. (b) qRT-PCR quantification of *RPS3*. Transcript levels of *RPS3* were normalized to *Cons4*, a constitutively expressed control gene. A distinct biological replicate independent of the samples used for RNA sequencing was used for relative transcript level (RTLs) quantifications.

**Figure S4. Identified co-expressed gene clusters in the four lines.** (a-d) All clusters grouped based on k-means clustering in Colfax (cf; a), NE2701 (ne; b), Senaki (sk; c), and Williams 82 (wm; d). The X-axis texts represent control (c) and inoculated (t) samples at different time points. The complete list of genes in these clusters can be found in Dataset S2.

**Figure S5. Proportion of transcribed TEs in each superfamily.** (a-j) Proportion of transcribed TEs (NCPK; normalized counts per kilobase of TE length > 0.5) in different lines and conditions. The proportions were averaged between the two replicates.

**Figure S6. Total transcript levels of TEs in each superfamily.** (a-j) The expression levels of TEs in ten superfamilies. The expression levels were determined by the mean normalized counts divided by the length of each TE (NCPK) for TEs within each superfamily, across different lines and conditions. The error bars represent SE.

**Figure S7. Transcription patterns of genes, lincRNAs and TEs.** (a) The proportion of transcribed transcripts for each transcript type. NCPK (normalized counts per kilobase of length of transcripts or genes) of all 64 samples was used for calculating the proportion of transcribed transcripts (NCPK > 0.5). (b) Comparison of transcript levels for four transcript types. Log(NCPK + 1) was used as a transcript level only for transcribed transcripts as depicted in (a). The red points indicate mean transcript levels. Different characters represent statistically significant differences between means (ANOVA followed by Tukey’s HSD test; *P* < 0.05).

**Figure S8. qRT-PCR quantification of *LOR1* (*Glyma.03G040900*).** Transcript levels of *LOR1* were normalized to *Cons4*, a constitutively expressed control gene. A distinct biological replicate independent of the samples used for RNA sequencing was used for transcript quantification.

**Figure S9. TE-containing lincRNAs exclusively differentially expressed in the resistant lines.** (a) The number of up-regulated TE-containing lincRNAs at 8 hpi. Among the different time points and treatments, only a single up-regulated lincRNA was found in common in the resistant lines. The red number represents the shared DE TE-containing lincRNA in the resistant lines. (b) Transcript levels of the DE TE-containing lincRNA, XLOC_059371, under different lines and conditions. Moreover, it was exclusively detected in the resistant lines at 8 hpi. A total of 49 co-expressed DEGs were linked with this lincRNA (|R| > 0.85; *P* < 0.001), but no adjacent DEG was found within 10 kb upstream or downstream. The TE within this lincRNA is classified as DNA/CACTA (ID: 294229).

**Figure S10. No significant changes in CG and CHG methylation on lincRNA bodies and their flanking regions upon pathogen inoculation.** (a) Mean proportions of CG and CHG methylation in non-TE lincRNAs for each line and condition. The error bars represent ± SE. Asterisks indicate statistically significant differences of means in inoculated plants compared to controls (Student’s *t* test; *P* ≥ 0.05, ns, not significant). (b) Distribution of CG and CHG methylation within 2 kb upstream and downstream of non-TE lincRNA transcript bodies. The mean methylation proportion was calculated in 40 windows for each of upstream, body, and downstream of transcripts. The lines were smoothed using LOESS.

**Table S1. Five most significantly enriched GO terms exclusive to each resistant line.**

**Table S2. Gene list in the enriched GO term, “response to ethylene” among co-expressed genes in the resistant lines (refer to Figure 3f).**

**Table S3. Summary of identified lncRNAs in soybean transcriptomes.**

**Table S4. Overrepresented GO terms of potential *tans*-target genes of DE lincRNAs (refer to Figure 5e).**

**Dataset S1. Description and normalized counts of DEGs upon inoculation in four soybean lines.**

**Dataset S2. Selected clusters for up-regulated genes upon inoculation.**

**Dataset S3. Enriched GO terms for genes in the selected clusters from Dataset S2.**

**Dataset S4. DE TEs upon inoculation and their location nearby or within DEGs or DE lincRNAs.**

**Dataset S5. DE lincRNAs upon inoculation with overlapping DE TEs.**

**Dataset S6. Correlated DEGs with DE non-TE lincRNAs as potential *trans*-targets (|R| > 0.85, *P* < 0.001).**

**Dataset S7. DEGs located within 5 kb upstream or downstream of DE non-TE lincRNAs as potential *cis*-targets.**

**Dataset S8. Correlated DEGs with the DE TE-containing lincRNA, XLOC_059371, exclusively in the resistant lines (|R| > 0.85, *P* < 0.001).**

## Notes

### Competing Interest Statement

The authors have declared no competing interest.

