## Supplemental Figures and Tables for "Transcriptomic and epigenetic responses shed light on soybean resistance to *Phytophthora sansomeana*"

**Supporting Information**

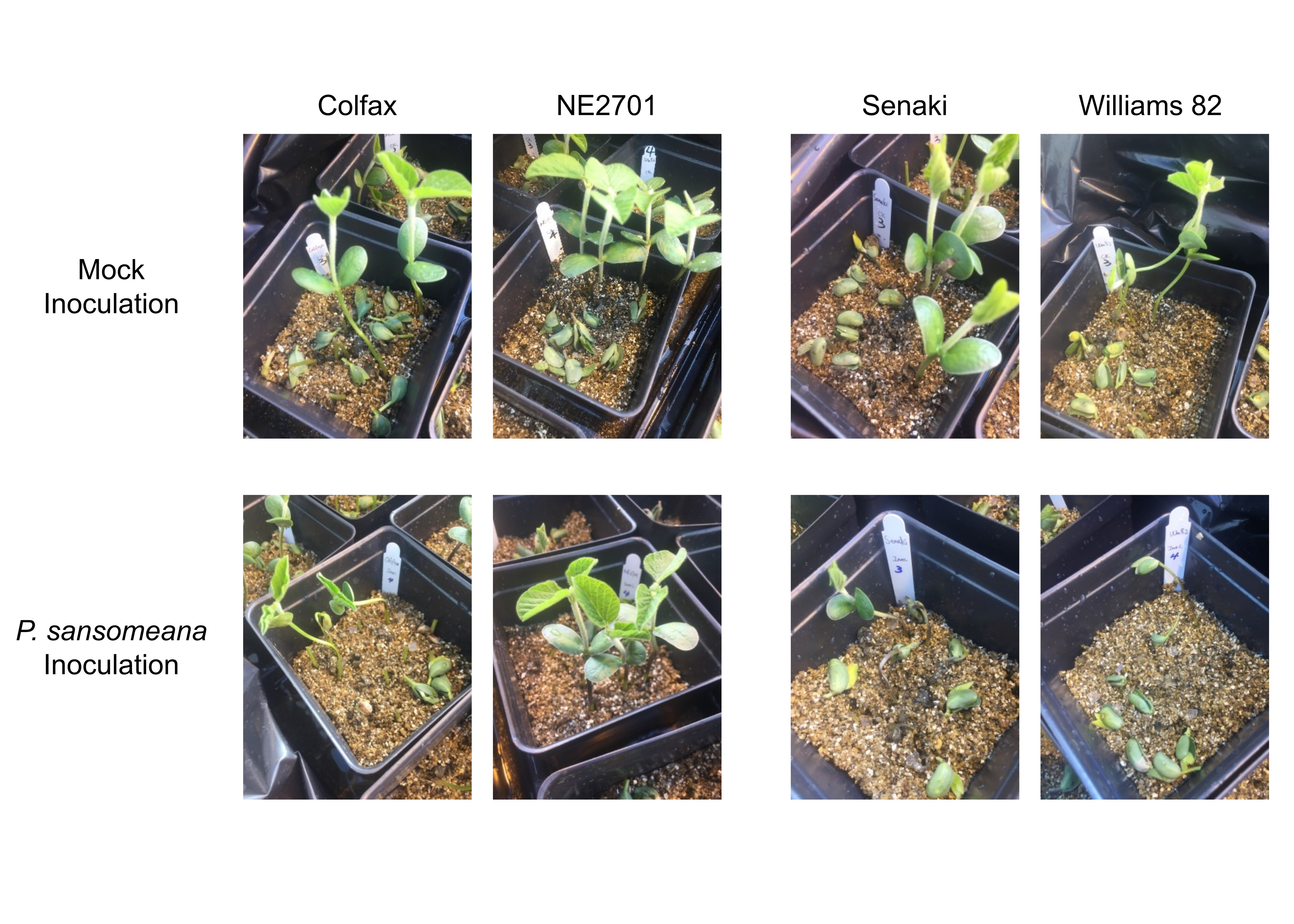

**Figure S1. Phenotypes of mock- and *P. sansomeana*-inoculated plants of the four soybean lines.** The photos were taken five days after mock- or pathogen-inoculation for four lines.

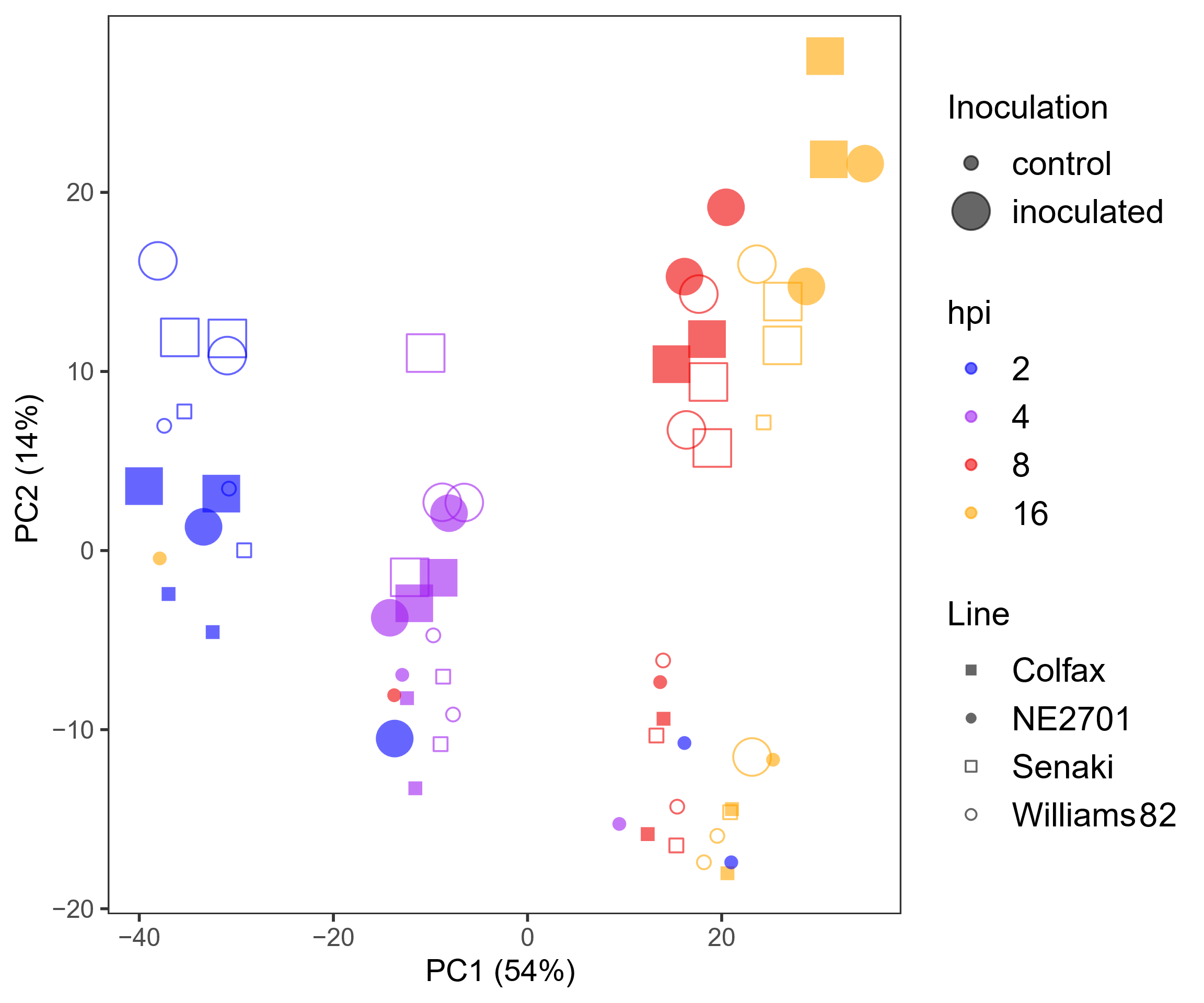

**Figure S2. Principal component analysis (PCA) of 64 soybean transcriptomes.** Variant stabilizing transformations of read counts were used for PCA to determine the sample distance in four soybean lines. The percentages of variation explained by PC1 and PC2 are indicated in parentheses.

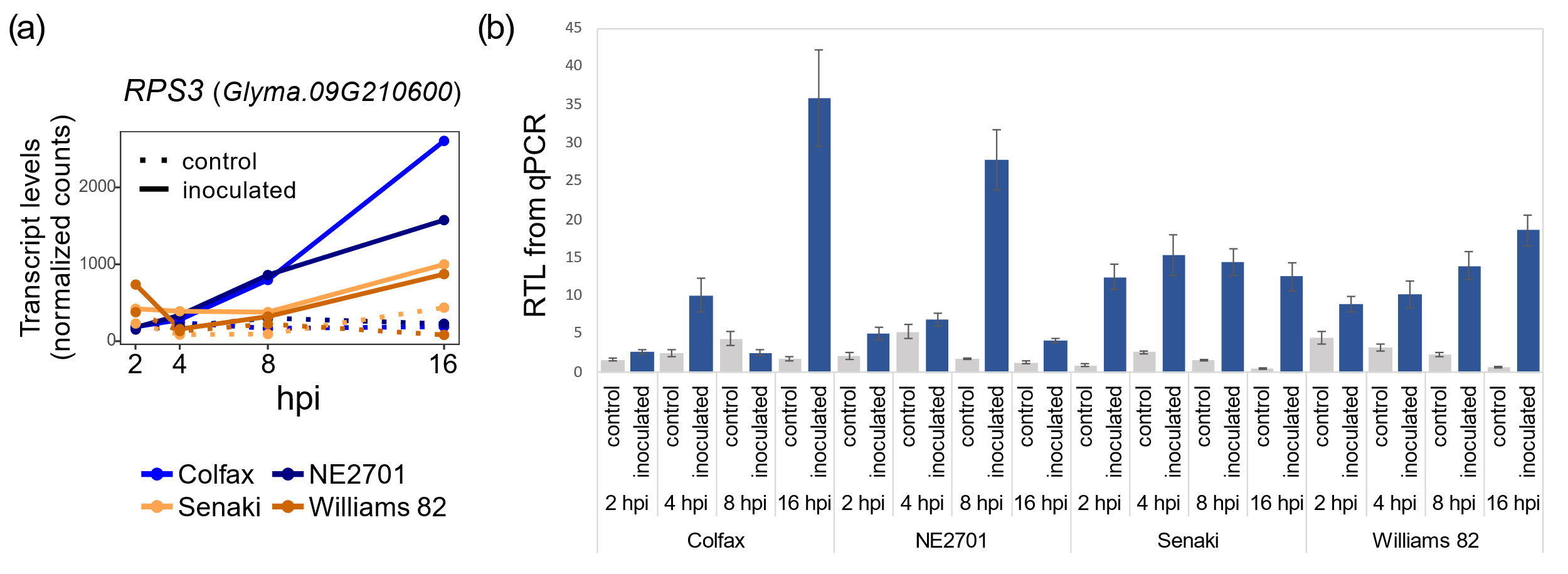

**Figure S3. Transcript levels of *Glyma.09G210600* (*RPS3*), exclusively up-regulated at 16 hpi in both resistant lines.** (a) Transcript levels of *RPS3* from RNA sequencing. Normalized counts were used for transcript levels at different time points for each line and condition. (b) qRT-PCR quantification of *RPS3*. Transcript levels of *RPS3* were normalized to *Cons4*, a constitutively expressed control gene. A distinct biological replicate independent of the samples used for RNA sequencing was used for relative transcript level (RTLs) quantifications.

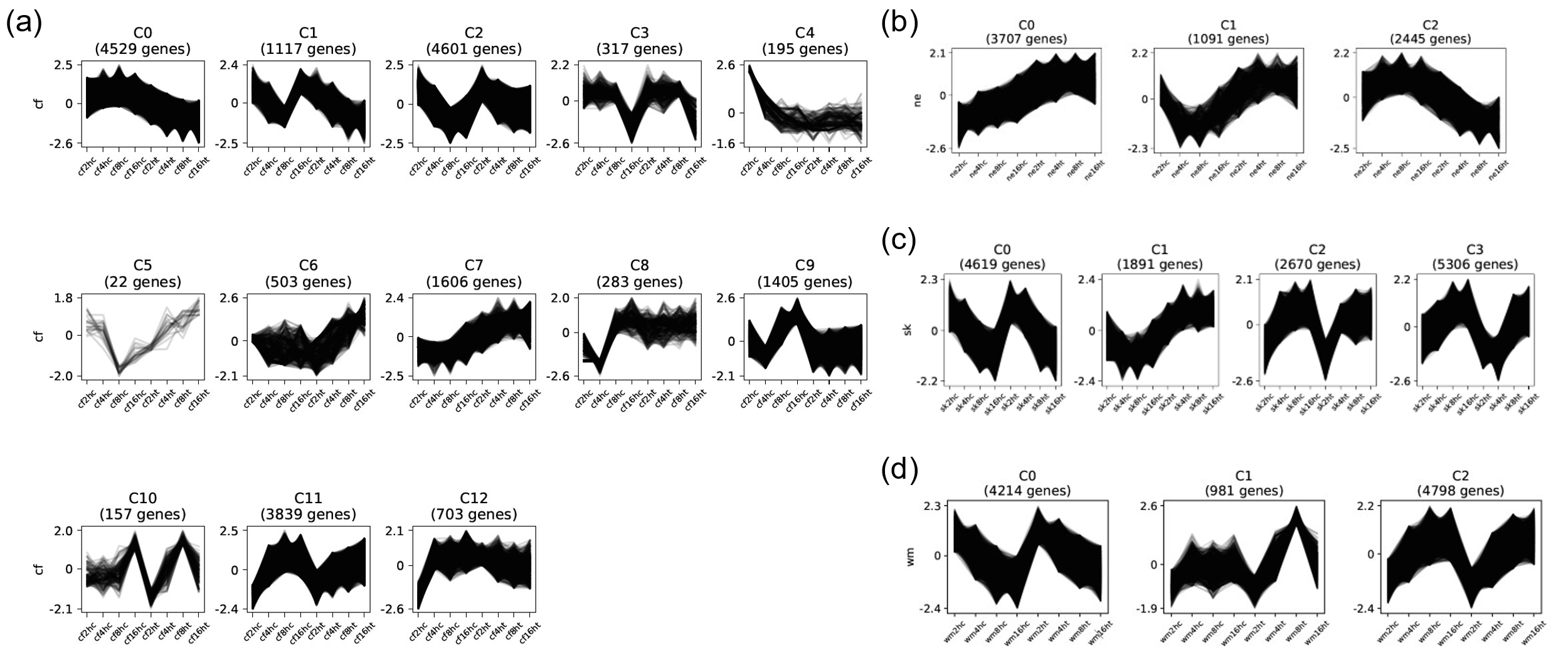

**Figure S4. Identified co-expressed gene clusters in the four lines.** (a-d) All clusters grouped based on k-means clustering in Colfax (cf; a), NE2701 (ne; b), Senaki (sk; c), and Williams 82 (wm; d). The X-axis texts represent control (c) and inoculated (t) samples at different time points. The complete list of genes in these clusters can be found in Dataset S2.

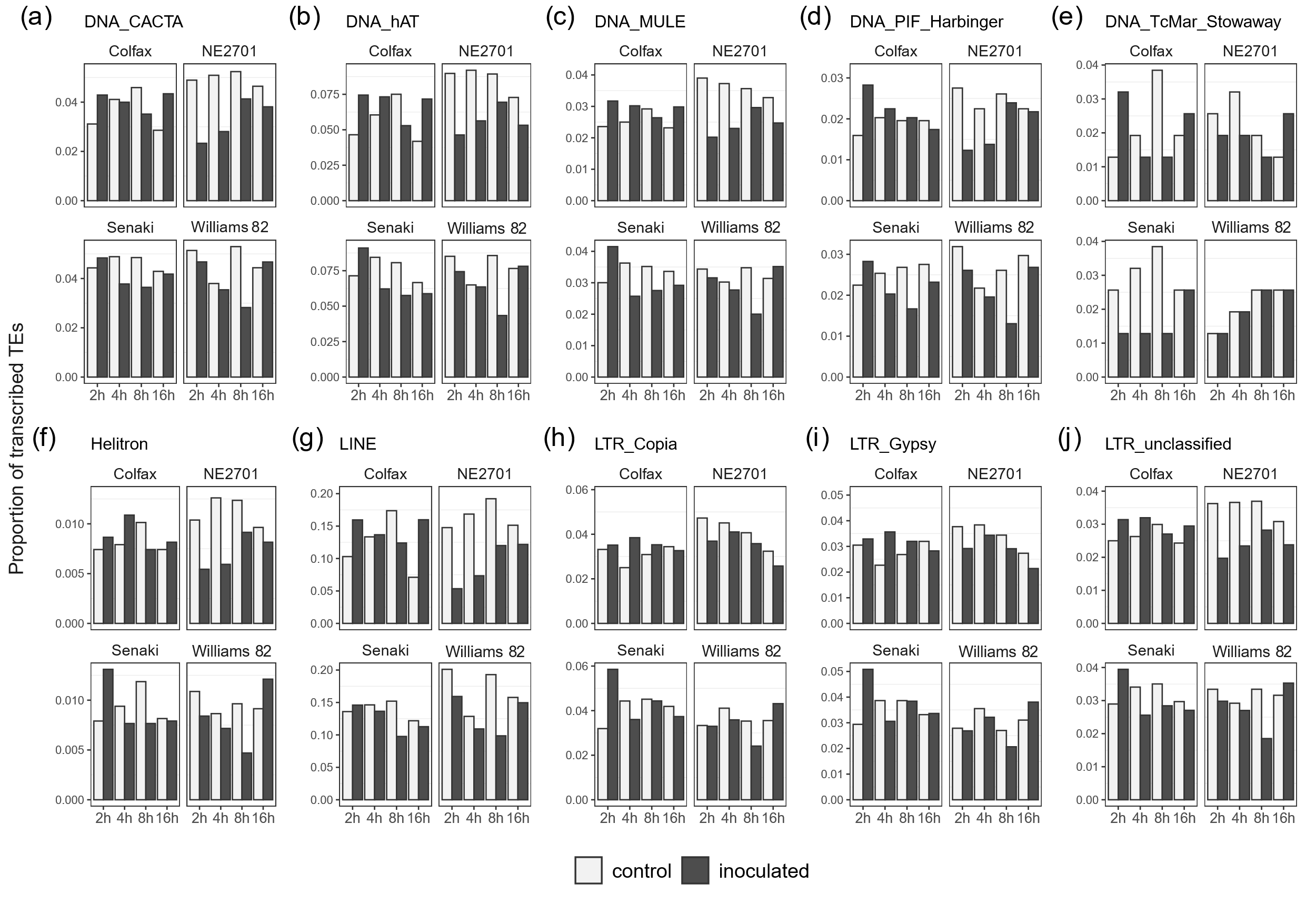

**Figure S5. Proportion of transcribed TEs in each superfamily.** (a-j) Proportion of transcribed TEs (NCPK; normalized counts per kilobase of TE length > 0.5) in different lines and conditions. The proportions were averaged between the two replicates.

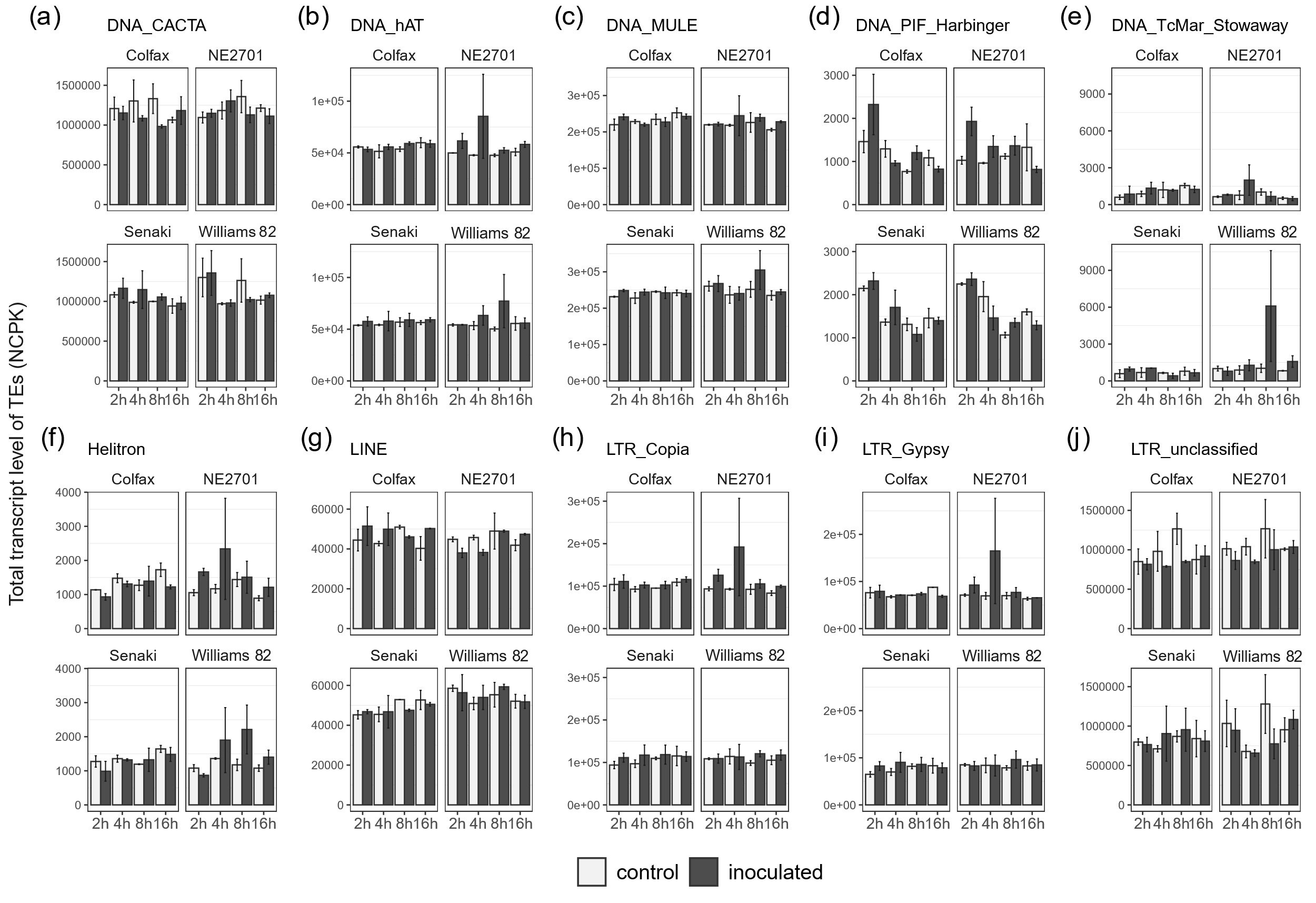

**Figure S6. Total transcript levels of TEs in each superfamily.** (a-j) The expression levels of TEs in ten superfamilies. The expression levels were determined by the mean normalized counts divided by the length of each TE (NCPK) for TEs within each superfamily, across different lines and conditions. The error bars represent SE.

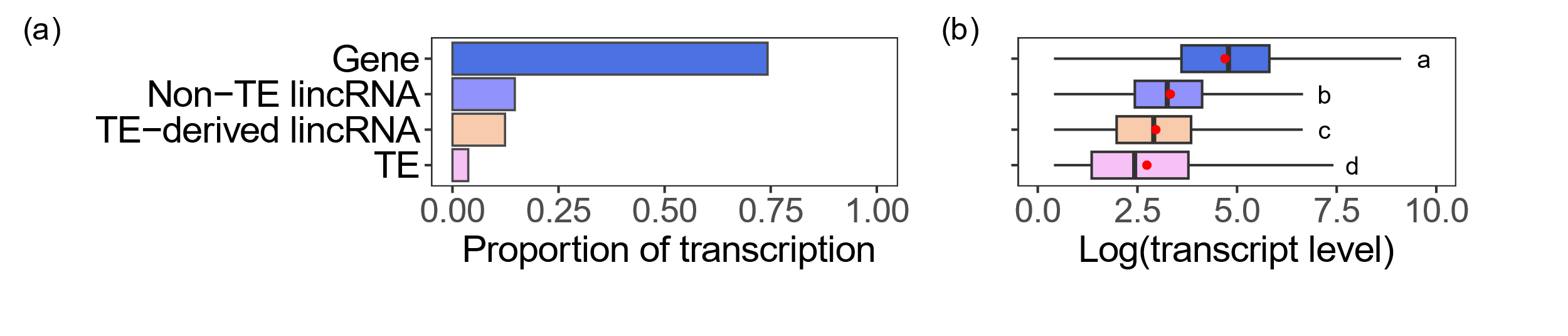

**Figure S7. Transcription patterns of genes, lincRNAs and TEs.** (a) The proportion of transcribed transcripts for each transcript type. NCPK (normalized counts per kilobase of length of transcripts or genes) of all 64 samples was used for calculating the proportion of transcribed transcripts (NCPK > 0.5). (b) Comparison of transcript levels for four transcript types. Log(NCPK + 1) was used as a transcript level only for transcribed transcripts as depicted in (a). The red points indicate mean transcript levels. Different characters represent statistically significant differences between means (ANOVA followed by Tukey’s HSD test; *P* < 0.05).

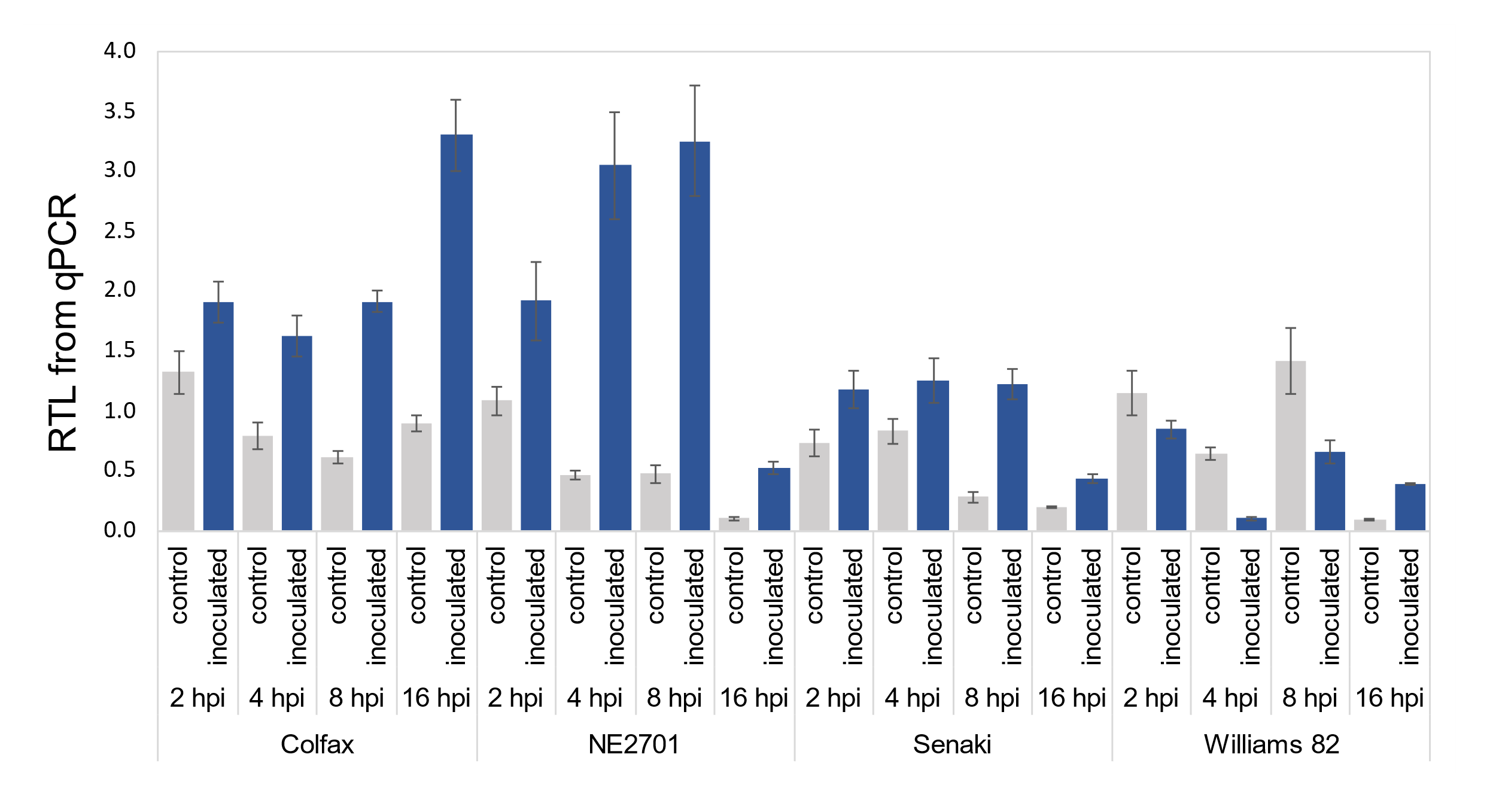

**Figure S8. qRT-PCR quantification of *LOR1* (*Glyma.03G040900*).** Transcript levels of *LOR1* were normalized to *Cons4*, a constitutively expressed control gene. A distinct biological replicate independent of the samples used for RNA sequencing was used for transcript quantification.

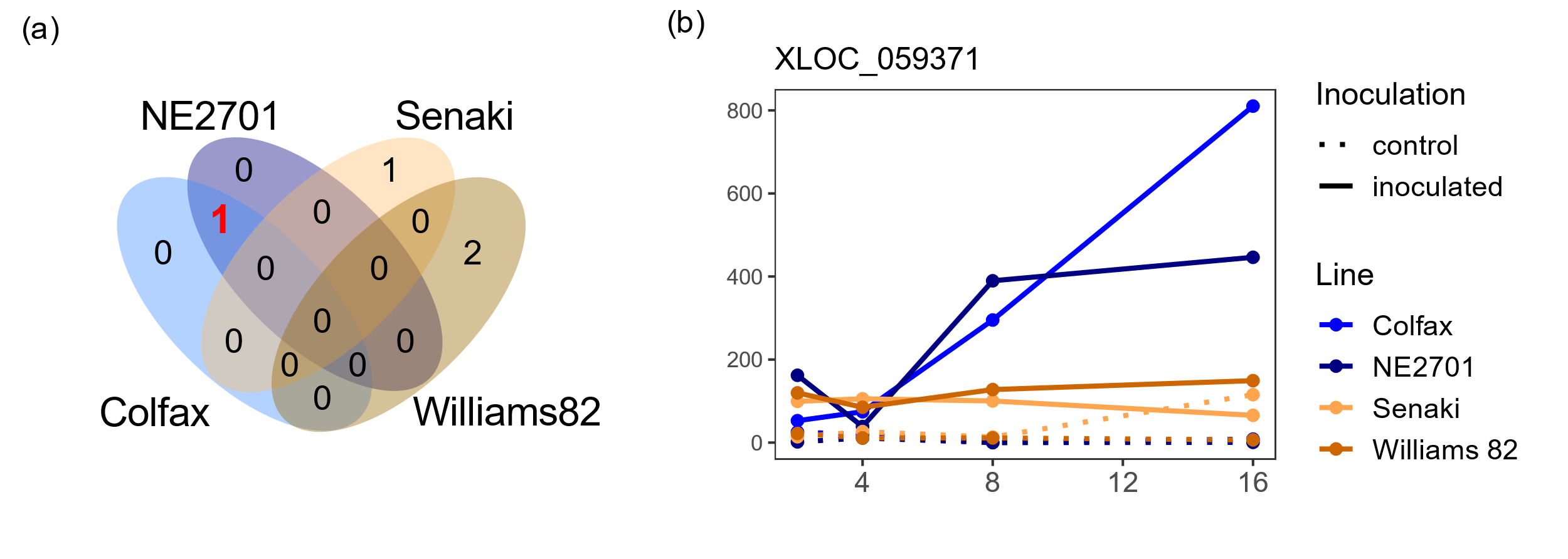

**Figure S9. TE-containing lincRNAs exclusively differentially expressed in the resistant lines.** (a) The number of up-regulated TE-containing lincRNAs at 8 hpi. Among the different time points and treatments, only a single up-regulated lincRNA was found in common in the resistant lines. The red number represents the shared DE TE-containing lincRNA in the resistant lines. (b) Transcript levels of the DE TE-containing lincRNA, XLOC_059371, under different lines and conditions. Moreover, it was exclusively detected in the resistant lines at 8 hpi. A total of 49 co-expressed DEGs were linked with this lincRNA (|R| > 0.85; *P* < 0.001), but no adjacent DEG was found within 10 kb upstream or downstream. The TE within this lincRNA is classified as DNA/CACTA (ID: 294229).

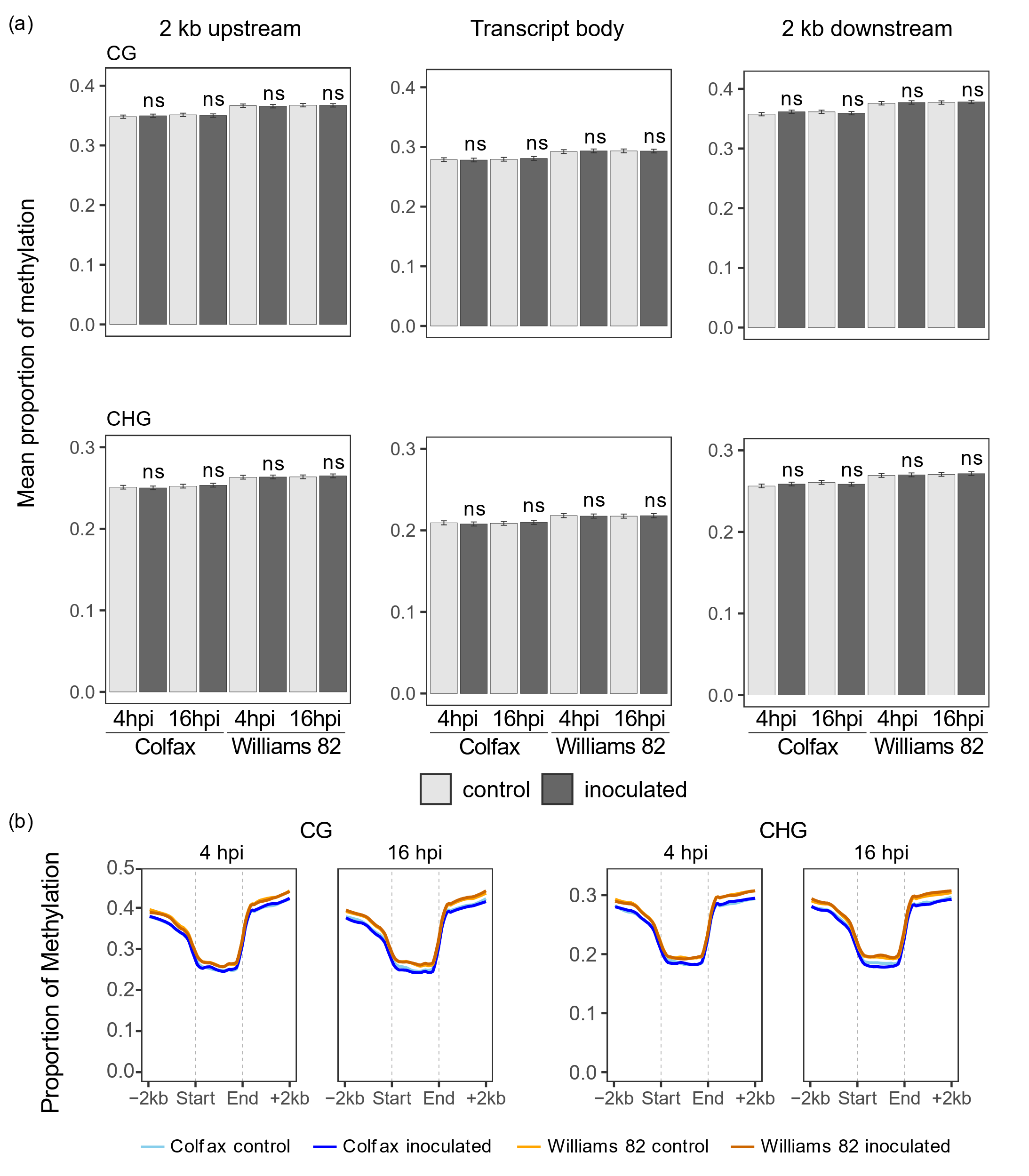
**Figure S10. No significant changes in CG and CHG methylation on lincRNA bodies and their flanking regions upon pathogen inoculation.** (a) Mean proportions of CG and CHG methylation in non-TE lincRNAs for each line and condition. The error bars represent ± SE. Asterisks indicate statistically significant differences of means in inoculated plants compared to controls (Student’s *t* test; *P* ≥ 0.05, ns, not significant). (b) Distribution of CG and CHG methylation within 2 kb upstream and downstream of non-TE lincRNA transcript bodies. The mean methylation proportion was calculated in 40 windows for each of upstream, body, and downstream of transcripts. The lines were smoothed using LOESS.

| Regulation | hpi^1^ | Line | GO term^2^ | adjusted *P* |
| --- | --- | --- | --- | --- |
| up | 2 | NE2701 | regulation of jasmonic acid mediated signaling pathway | 3.12E-15 |
|  |  |  | cellular response to fatty acid | 1.12E-11 |
|  |  |  | response to fatty acid | 1.58E-11 |
|  |  |  | response to wounding | 5.50E-11 |
|  |  |  | cellular response to jasmonic acid stimulus | 8.50E-11 |
| down | 2 | NE2701 | plant-type cell wall organization | 1.95E-05 |
|  |  |  | plant-type cell wall organization or biogenesis | 2.40E-05 |
|  |  |  | anatomical structure morphogenesis | 2.90E-05 |
|  |  |  | cell wall organization or biogenesis | 9.19E-05 |
|  |  |  | cellular response to chemical stimulus | 9.25E-05 |
| up | 16 | Colfax | phosphate-containing compound metabolic process | 1.68E-21 |
|  |  |  | phosphorus metabolic process | 3.10E-20 |
|  |  |  | oxoacid metabolic process | 6.58E-20 |
|  |  |  | phosphorylation | 1.15E-18 |
|  |  |  | organic acid metabolic process | 4.84E-18 |
| down | 16 | Colfax | photosynthesis | 1.28E-43 |
|  |  |  | photosynthesis, light reaction | 5.98E-23 |
|  |  |  | photosynthesis, light harvesting in photosystem I | 6.74E-18 |
|  |  |  | photosynthesis, light harvesting | 8.49E-17 |
|  |  |  | nucleosome assembly | 1.82E-15 |

**Table S1. Five most significantly enriched GO terms exclusive to each resistant line.**

^1^The time points were selected based on hpi at which the greatest number of exclusive DEGs in each of resistant lines was found.

^2^GO terms enriched as biological process.

**Table S2. Gene list in the enriched GO term, “response to ethylene” among co-expressed genes in the resistant lines (refer to Figure 3f).**

| Gene ID | Name | Description | Top Arabidopsis BLASTP Hit |
| --- | --- | --- | --- |
| Glyma.10G036700 | *ERF1* | ethylene response factor | AT3G23240 |
| Glyma.13G122700 | *ERF1* | ethylene response factor | AT3G23240 |
| Glyma.03G162500 | *ERF15* | ethylene response factor | AT2G31230 |
| Glyma.08G169100 | *S8H* | 2-oxoglutarate (2OG) and Fe(II)-dependent oxygenase superfamily protein | AT3G12900 |
| Glyma.09G052800 | *ERF98* | ethylene response factor | AT3G23230 |
| Glyma.10G036600 | *ERF98* | ethylene response factor | AT3G23230 |
| Glyma.14G049500 | *ACO4* | ethylene-forming enzyme | AT1G05010 |
| Glyma.15G159200 | *ERF98* | ethylene response factor | AT3G23230 |
| Glyma.19G164000 | *ERF98* | ethylene response factor | AT3G23230 |

**Table S3. Summary of identified lncRNAs in soybean transcriptomes.**

|  | Non-TE  lincRNA^1^ | TE-containing lincRNA | SOT^2^ | AOT^3^ |
| --- | --- | --- | --- | --- |
| Total number of lncRNAs | 13,487 | 27,672 | 1,714 | 886 |
| Total length (bp) | 6,259,204 | 11,180,024 | 2,437,046 | 988,707 |
| Smallest lincRNA (bp) | 201 | 201 | 201 | 201 |
| Largest lincRNA (bp) | 8,558 | 7,664 | 8,287 | 8,086 |
| Average length of lincRNA (bp) | 464 | 404 | 1,422 | 1,116 |

^1^Long intergenic non-coding RNAs that have no TEs

^2^Sense-overlapping lncRNA transcripts

^3^Antisense-overlapping lncRNA transcripts

| LincRNA | Number of trans-targets | GO category | GO term name | GO term ID | adjusted *P* |
| --- | --- | --- | --- | --- | --- |
| XLOC_014629 | 101 | GO:MF | catalytic activity | GO:0003824 | 0.0019 |
|  |  | GO:MF | FMN reductase (NADPH) activity | GO:0052873 | 0.0075 |
|  |  | GO:BP | secondary metabolic process | GO:0019748 | 0.0009 |
|  |  | GO:BP | phenylpropanoid biosynthetic process | GO:0009699 | 0.0026 |
|  |  | GO:BP | isopentenyl diphosphate biosynthetic process | GO:0009240 | 0.0029 |
|  |  | GO:BP | isopentenyl diphosphate metabolic process | GO:0046490 | 0.0029 |
|  |  | GO:BP | secondary metabolite biosynthetic process | GO:0044550 | 0.0083 |
|  |  | KEGG | Isoflavonoid biosynthesis | KEGG:00943 | 0.0039 |
| XLOC_084885 | 75 | GO:MF | oxidoreductase activity | GO:0016491 | 0.0017 |
|  |  | GO:MF | catalytic activity | GO:0003824 | 0.0050 |
|  |  | GO:BP | secondary metabolic process | GO:0019748 | 0.0001 |
|  |  | GO:BP | phenylpropanoid biosynthetic process | GO:0009699 | 0.0005 |
|  |  | GO:BP | isopentenyl diphosphate biosynthetic process | GO:0009240 | 0.0010 |
|  |  | GO:BP | isopentenyl diphosphate metabolic process | GO:0046490 | 0.0010 |
|  |  | GO:BP | secondary metabolite biosynthetic process | GO:0044550 | 0.0016 |
|  |  | GO:BP | phenylpropanoid metabolic process | GO:0009698 | 0.0033 |
|  |  | GO:BP | isoprenoid biosynthetic process | GO:0008299 | 0.0037 |
|  |  | GO:BP | cellular metabolic process | GO:0044237 | 0.0039 |
|  |  | GO:BP | lignan metabolic process | GO:0009806 | 0.0048 |
|  |  | GO:BP | lignan biosynthetic process | GO:0009807 | 0.0048 |
| XLOC_063809 | 26 | GO:BP | salicylic acid biosynthetic process | GO:0009697 | 0.0000 |
|  |  | GO:BP | regulation of salicylic acid metabolic process | GO:0010337 | 0.0000 |
|  |  | GO:BP | regulation of salicylic acid biosynthetic process | GO:0080142 | 0.0000 |
|  |  | GO:BP | phenol-containing compound biosynthetic process | GO:0046189 | 0.0000 |
|  |  | GO:BP | regulation of cellular ketone metabolic process | GO:0010565 | 0.0002 |
|  |  | GO:BP | salicylic acid metabolic process | GO:0009696 | 0.0004 |
|  |  | GO:BP | regulation of small molecule metabolic process | GO:0062012 | 0.0005 |
|  |  | GO:BP | phenol-containing compound metabolic process | GO:0018958 | 0.0006 |
|  |  | GO:BP | benzene-containing compound metabolic process | GO:0042537 | 0.0023 |

**Table S4. Overrepresented GO terms of potential *tans*-target genes of DE lincRNAs (refer to Figure 5e).**
